# Public-Data Reanalysis Links MS4A4A to M2-like Human Myeloid States and Supports a Predicted Four-Pass Transmembrane Fold

**DOI:** 10.64898/2026.08.19.745820

**Authors:** Shi-Ruei Lin, Mandarina Li, Sihan Wang, Eric Li, Hao Sun, Liangyu Li

**Affiliations:** South Hills Academy, 1600 E Francisquito Ave, West Covina, CA 91791, USA; Wolfson Institute for Biomedical Research, Division of Medicine, University College London, Gower Street, London WC1E 6BT, United Kingdom; Faculty of Information Science and Technology, Universiti Kebangsaan Malaysia, Bangi 43600, Selangor, Malaysia; Faculty of Economics and Business, Universidad Complutense de Madrid, Campus de Somosaguas, 28223 Pozuelo de Alarcón, Madrid, Spain

**Keywords:** . MS4A4A, MS4A6A, macrophage polarization, M2 macrophage, chromatin accessibility, single-cell RNA-seq, AlphaFold, TREM2

## Abstract

**Background/Objectives:** MS4A4A is associated with M2-like macrophage states, while the MS4A gene cluster modifies soluble TREM2 levels and Alzheimer’s disease risk. We asked whether MS4A4A consistently marks the M2 side of human myeloid activation and whether AlphaFold supports a proposed MS4A4A–MS4A6A interaction. Methods: We reanalysed four public human datasets: bulk RNA-seq and ATAC-seq of primary monocyte-derived macrophages from independent three-donor cohorts, and single-cell RNA-seq atlases of healthy liver and severe COVID-19 blood. MS4A4A and an MS4A4A–

MS4A6A complex were modelled with AlphaFold 3 and evaluated using pLDDT, predicted aligned error, and ipTM. Results: MS4A4A was higher in M2 (IL-4) than in M1 (IFN-γ + LPS) macrophages in all three donors (log2 fold change +2.68, adjusted P = 0.0013). Its promoter showed the highest mean accessibility in M2. MS4A4A was macrophage-enriched in liver and monocyte-enriched in blood, and was detected in 76.7% of M2-like versus 39.3% of M1-like liver macrophages, with the difference driven mainly by the proportion of positive cells. AlphaFold confidently modelled the four transmembrane helices (mean pLDDT 83.1), but the predicted MS4A4A–MS4A6A interface was not supported (ipTM 0.59). Conclusions: MS4A4A is consistently associated with the M2 side of human myeloid activation across independent transcriptomic, chromatin, and single-cell datasets. The findings are associative, and the proposed MS4A4A–MS4A6A interface remains an untested structural hypothesis.

## 1. Introduction

Macrophages are highly plastic innate immune cells with essential roles in tissue homeostasis, host defence, inflammation, and repair. In response to environmental and cellular signals they adopt distinct functional states with different transcriptional and immunological programmes.

Macrophage activation is a continuum rather than a pair of discrete populations, but the M1/M2 framework remains a useful description of its two best characterized endpoints: M1-like states, induced experimentally by interferon-γ (IFN-γ) and lipopolysaccharide (LPS), are associated with pro-inflammatory responses and host defence [1,2], whereas M2-like states, induced by interleukin 4 (IL-4), are associated with tissue repair, immune regulation, and the resolution of inflammation [1,3]. Throughout this paper we use "M1" and "M2 (IL-4)" for cells polarized in culture and "M1-like" and "M2-like" for cells annotated post hoc in tissue, and we treat these as endpoints on a spectrum rather than as fixed cell types. Following convention, gene symbols are set in italic and protein names in roman. Dysregulation of macrophage states is implicated in chronic inflammatory disease and in cancer, so molecules that distinguish macrophage functional states are of interest both as markers and as potential points of intervention [4].

One such candidate is membrane spanning 4 domains subfamily A member 4A (MS4A4A), a member of the MS4A family of tetraspan membrane proteins [5]. *MS4A4A* is expressed predominantly in the monocyte and macrophage lineage; its expression increases as monocytes differentiate into macrophages and is induced by IL-4 and by glucocorticoids [6,7]. In human tissue it has been detected in tissue-resident macrophages and in tumour-associated macrophages [5,8], and in macrophages associated with inflammatory disease [7]. MS4A4A is not only a marker: it enhances Dectin-1 dependent signalling and natural killer cell mediated resistance to metastasis [8], and reducing its expression alters M2-associated markers and macrophage behaviour in glioblastoma models [9]. These observations suggest MS4A4A participates in macrophage biology rather than merely reflecting activation state.

The association between *MS4A4A* and M2-like programmes matters clinically because such programmes can sustain inflammation and shape the tumour microenvironment, where immunoregulatory and tissue remodelling macrophages can support tumour progression and blunt anti tumour immunity [5,9]. Interest in MS4A4A has grown further because of its connection to TREM2 signalling. TREM2 is a myeloid receptor that senses lipids and cellular damage and regulates phagocytosis, survival, and inflammatory responses [10,11]. Common variants across the *MS4A* gene cluster are among the strongest known modifiers of soluble TREM2 concentration and of Alzheimer’s disease risk [12], and MS4A4A and MS4A6A have been shown to cooperate to negatively regulate TREM2 through DAP12, with experimental reduction of MS4A4A altering TREM2 abundance and microglial state [13]. This gives the MS4A family a mechanistic rationale as a therapeutic target rather than a biomarker alone, and antibody-based approaches directed at MS4A are being explored clinically: a Phase 1 study of AL044, an antibody designed to phenocopy protective *MS4A* variants and raise TREM2 levels, has been announced by its sponsor, though the results are not yet peer reviewed [14].

Despite this functional evidence, a structural gap remains. MS4A4A has four predicted transmembrane domains, but no experimentally determined three dimensional structure of the protein is available; the only MS4A family member with a solved structure is CD20 (MS4A1), determined by cryo-electron microscopy as a homodimer in complex with therapeutic antibody Fab fragments [15]. The absence of an MS4A4A structure limits understanding of how its transmembrane helices are organized and how the protein might interact with other membrane proteins, a question made concrete by the reported functional partnership with MS4A6A. Structure prediction offers one route to a testable hypothesis: AlphaFold and related methods generate models that are particularly useful for membrane proteins, which resist experimental structure determination [16–18], provided the models are read as hypotheses rather than as evidence, and provided their confidence measures are reported. For multi chain predictions this last point is decisive, because a model can be confident about each chain and simultaneously uninformative about how the chains are arranged.

In this study we characterized MS4A4A from both a biological and a structural perspective, using only public data (Table S1). We first reanalysed bulk RNA sequencing (RNA-seq) and bulk assay for transposase accessible chromatin with sequencing (ATAC-seq) [19] of primary human monocyte-derived macrophages to ask whether *MS4A4A* expression and promoter accessibility are elevated in the M2 state. We then used two single-cell RNA-seq atlases, one of healthy human liver and one of peripheral blood from patients with severe coronavirus disease 2019 (COVID-19) and healthy donors, to ask whether the association survives at single-cell resolution and across cell types. Finally, we used AlphaFold to predict the structure of the MS4A4A monomer and of a candidate MS4A4A-MS4A6A complex, and evaluated both with per-residue confidence (predicted local distance difference test, pLDDT), predicted aligned error (PAE), and, for the complex, the interface predicted template modelling score (ipTM) [20], which is the diagnostic the preceding paragraph identifies as decisive. Together these analyses test whether *MS4A4A* is a reproducible molecular feature of the M2 side of the human myeloid activation range, and provide a structural framework, with its uncertainty made explicit, for future experimental work on MS4A4A and MS4A6A.

## 2. Materials and Methods

### 2.1. Data Sources

All data analysed in this study are previously published and publicly available; no new experimental data were generated. The four primary datasets and the two reference structures are listed in Table S1. Both single-cell datasets were retrieved through CZ CELLxGENE Discover (liver dataset a43aa46b-bd16-47fe-bc3e-19a052624e79; blood dataset 456e8b9b- f872-488b-871d-94534090a865) [21]; the Gene Expression Omnibus (GEO) accessions in Table S1 are the primary sources and are what we cite. The input matrices for every analysis reported here, together with all analysis scripts and the data provenance records that describe them, are supplied as Supplementary Software S1.

### 2.2. Bulk RNA-seq Differential Expression

#### 2.2.1. Dataset and Count Matrix

GSE162698 comprises primary human monocyte-derived macrophages from three donors, polarized for 24 h to M0 (no stimulus), M1 (20 ng/mL IFN-γ + 100 ng/mL LPS), M2/IL-10 (20 ng/mL IL-10), M2/IL-4 (20 ng/mL IL-4), or a tumour-associated macrophage (TAM)-like state (A549 tumour conditioned medium + 20 ng/mL macrophage colony stimulating factor); 15 samples in total [22]. We retained the 12 samples corresponding to M0, M1, M2/IL-4, and TAM-like. The M2/IL-10 arm was excluded so that "M2" refers unambiguously to the IL-4 repair state throughout this paper.

GEO provides gene-level transcripts per million (TPM) for this series, not raw counts. Because DESeq2 requires non negative integers, each TPM value *t* was converted to a pseudocount as round(*t* × 20); the factor of 20 scales a per-sample TPM total of 10^6^ to approximately 2 × 10^7^, a realistic sequencing depth. This is a monotonic per-sample rescaling that preserves relative expression within a sample but does not recover true count level variance (see Section 4.4).

Genes were retained if their pseudocount was at least 10 in at least 3 of the 12 samples, and were keyed by HGNC symbol, keeping the first occurrence where several Ensembl identifiers mapped to one symbol. The final matrix comprised 12,900 genes by 12 samples.

#### 2.2.2. Differential Expression Testing

Testing was performed with DESeq2 1.52.0 in R 4.6.1 [23]. The model was ‘∼ condition + donor’, treating donor as a blocking factor to reflect the paired design, and the reported contrast is M2 versus M1, so that a positive log_2_ fold change indicates higher expression in M2. *P* values were adjusted across genes by the Benjamini and Hochberg false discovery rate procedure [24] as implemented in DESeq2. Genes were called differentially expressed at adjusted *P* < 0.05 and absolute log_2_ fold change > 1. The complete results table is provided as Table S2.

#### 2.2.3. Volcano Plot and Per Donor Plot (Figure 1a,b)

The volcano plot in Figure 1a was drawn from the results table in Table S2, following the four category colouring of EnhancedVolcano 1.30.0 [25], plotting the negative log_10_ of the adjusted *P* value against log_2_ fold change with thresholds at adjusted *P* = 0.05 and absolute log_2_ fold change = 1; *MS4A4A* was overplotted as a labelled point. The x axis is limited to plus and minus 16.5, which contains every gene called significant; five transcripts fall outside it (*MIR659*, *MIR550A1*, *MIR106B*, *RNU6ATAC*, and *SNORD71*), none of which is significant on either adjusted or nominal *P*.

**Figure 1.**
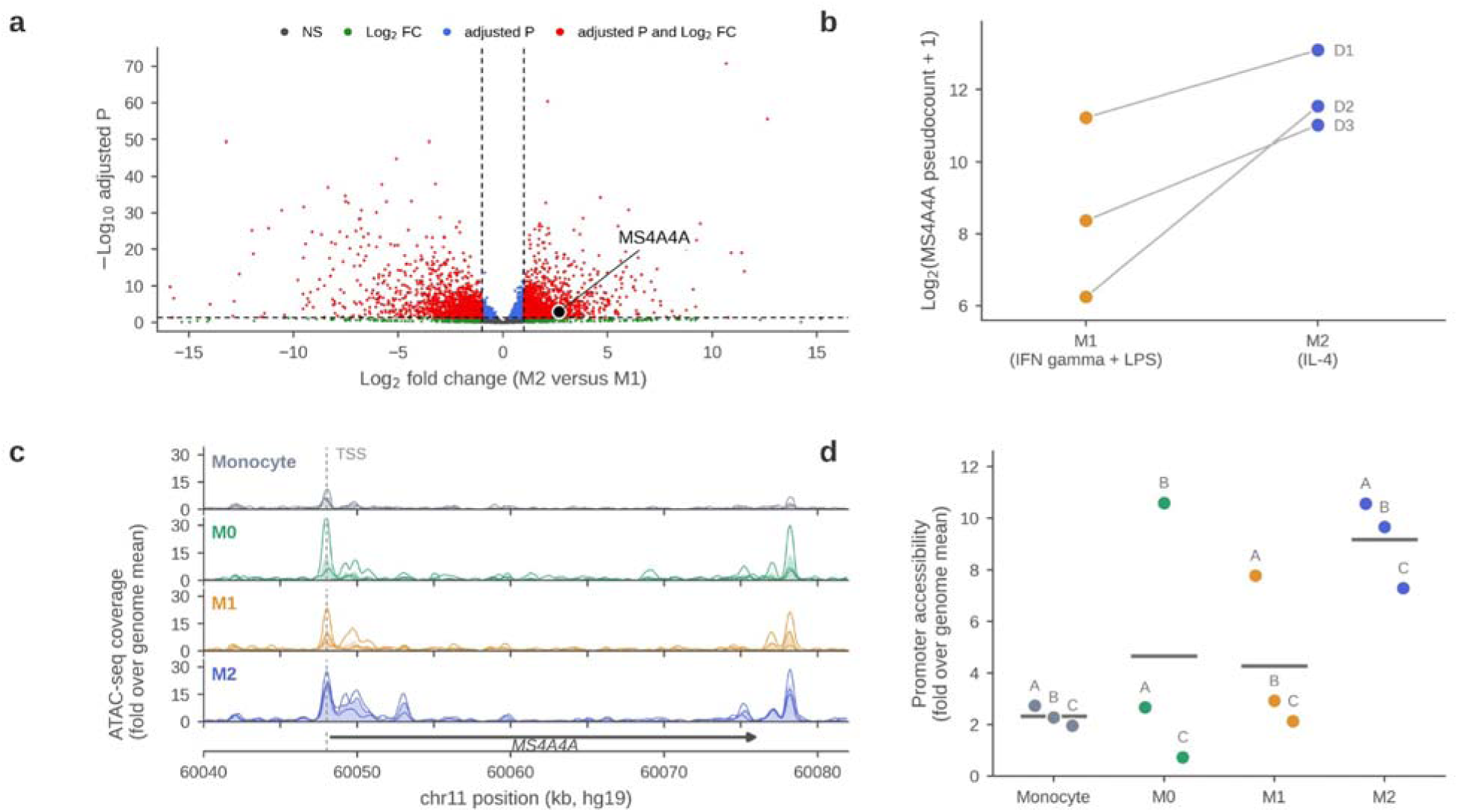
MS4A4A is transcriptionally upregulated and its promoter becomes more accessible in M2 macrophages. **(a)** Volcano plot of differential gene expression between M2 (IL-4) and M1 (IFN-γ + LPS) primary human monocyte-derived macrophages (GEO GSE162698; 12,900 genes; three donors; DESeq2, design ‘∼ condition + donor’), drawn from the results table in Table S2. Positive log2 fold change indicates higher expression in M2. Dashed lines mark the thresholds used to call differential expression (adjusted P < 0.05, absolute log2 fold change > 1); 1,534 genes were higher in M2 and 1,692 higher in M1. MS4A4A (black point, white outline) passes both thresholds (log2 fold change +2.68, adjusted P = 0.0013). The x axis spans plus and minus 16.5, which contains every significant gene; five non-significant small RNA transcripts lie outside it. **(b)** MS4A4A expression in matched M1 and M2 macrophages from the same three donors (GSE162698), as log2(pseudocount + 1). Each point is one donor, labelled D1 to D3; lines join matched samples from the same donor. **(c)** ATAC-seq coverage across the MS4A4A locus (hg19 chr11:60,040,000 to 60,082,000) in primary human monocytes and in M0, M1, and M2 (IL-4) macrophages from three donors (GEO GSE261697). Thin lines are individual donors; shaded areas are the donor mean. Coverage is normalized to each sample’s genome-wide mean, and is lightly smoothed for display only. The dashed line marks the MS4A4A promoter (transcription start site, chr11:60,048,014); the bar below shows the gene body and direction of transcription. **(d)** Quantification of (c): MS4A4A promoter accessibility, calculated as mean normalized coverage over chr11:60,047,000 to 60,049,599, for each donor separately. Each point is one donor (A to C); horizontal bars are the group mean. Expression values in (a,b) are integer pseudocounts derived from published TPM (see Section 2.2.1); the direction of the MS4A4A effect is robust, but the exact P values are approximate. Accessibility data in (c,d) derive directly from public coverage files and were normalized as described in Section 2.3.2; coverage completeness per sample is reported in Section 2.3.4.

For Figure 1b, *MS4A4A* pseudocounts were transformed as log_2_(*x* + 1) and plotted for the M1 and M2 sample of each donor as individual points, with lines connecting the paired samples from the same donor. No summary box or bar is drawn, because with three donors the paired lines carry the whole of the information. The underlying per-donor values are given in Table S3.

#### 2.2.4. Model Cross Check

To assess how much the paired design contributed, the same M2 versus M1 contrast was retested without donor blocking (’∼ condition’) using an independent implementation, pydeseq2 0.5.2 [26]. This gave *MS4A4A* log_2_ fold change +2.33 (adjusted *P* = 0.027), against +2.68 (adjusted *P* = 0.0013) with donor blocking, the same direction and a similar effect size. Note that this comparison changes the implementation as well as the design, because ‘∼ condition’ was not also run in DESeq2; it is therefore a robustness check rather than an isolated test of donor blocking. The unblocked results table is provided as Table S10.

### 2.3. Bulk ATAC-seq Chromatin Accessibility

#### 2.3.1. Dataset

GSE261697 provides ATAC-seq of primary human monocytes and monocyte-derived macrophages. The series as deposited in GEO contains 17 samples spanning primary, bone marrow-derived, and induced pluripotent stem cell derived monocytes and macrophages; we used the 12 samples comprising three donors (A, B, C) in four states: circulating monocyte, M0 (macrophage colony stimulating factor), M1 (IFN-γ + LPS), and M2 (macrophage colony stimulating factor followed by IL-4). Coverage was supplied as per-sample bigWig files aligned to GRCh37/hg19. No publication is linked to this accession; it is cited as a dataset [27].

#### 2.3.2. Coverage Extraction and Normalization

Signal was extracted across the MS4A4A locus, hg19 chr11:60,040,000 to 60,082,000 (42 kb), in bins of 100 bp (420 bins per-sample; 5,040 rows in total). The full bigWig files, up to 124 MB each, exceeded the available transfer limit, so extraction used a dependency free bigWig reader written for this study in Python 3 (standard library ‘struct’ and ‘zlib’ only; ‘bwslice.py’, provided in Supplementary Software S1), which parses the file header, R-tree index, and only those compressed data blocks overlapping the query window. The reader was validated against pyBigWig, reproducing its values exactly at the MS4A4A promoter, at GAPDH, at ACTB, and for the genome-wide mean. Extracted values were cached locally and are provided as Table S11. Each sample’s coverage was then divided by its genome-wide mean, taken from the total summary record embedded in its bigWig file, giving fold over genome mean values comparable across samples and donors.

#### 2.3.3. Coverage Tracks and Promoter Quantification (Figure 1c,d)

For Figure 1c, the three donor tracks of each state are plotted as thin lines with the donor mean as a shaded area; tracks were lightly smoothed with a five bin moving average for display only, and all reported values are computed from unsmoothed coverage. A dashed line marks the *MS4A4A* transcription start site (chr11:60,048,014); the gene body (chr11:60,048,014 to 60,076,445, + strand) is drawn beneath. Accessibility peaks were located as local maxima of the donor mean track exceeding 2 fold over genome mean; two intragenic peaks (chr11:60,049,900 and chr11:60,053,000) and one peak immediately downstream of the gene (chr11:60,078,200) are referred to in the Results.

For Figure 1d, promoter accessibility was quantified per-donor as the mean normalized coverage over chr11:60,047,000 to 60,049,599, a 2.6 kb window (26 bins of 100 bp) spanning 1.0 kb upstream to 1.6 kb downstream of the transcription start site. The window was chosen to contain the core promoter peak while excluding the intragenic peak at 60,049,900. Points are individual donors and horizontal bars are group means. Values are given in Table S6.

#### 2.3.4. Quality Control

No depth or alignment quality metrics are deposited with GSE261697, and the source bigWig files were not retained after extraction, so coverage completeness within the extracted window is the only quality measure we can report. We report it because the samples differ substantially, and because two of them carry the low values that the reproducibility comparison in the Results depends on. Across the 420 bins spanning the locus, the fraction of bins with exactly zero coverage was 63.1% in the donor A monocyte sample and 54.5% in the donor C M0 sample, against 18.8% to 42.6% in the other two M0 samples and the other two monocyte samples, 23.6% to 27.6% in the three M1 samples, and 21.9% to 24.5% in the three M2 samples. Within the 26 bin promoter window the pattern is sharper: all three M2 samples have coverage in every bin, the three M1 samples each lack coverage in 2 of 26 bins, and the two extreme samples lack 16 of 26 (donor C, M0, including the bin at the promoter apex) and 12 of 26 (donor A, monocyte). The values reported for those two samples should therefore be read as lower bounds, and the M2 versus M0 reproducibility comparison is correspondingly weaker than the M2 versus M1 comparison. An earlier draft of this work reported *GAPDH* promoter accessibility as a normalization check; those values could not be traced to any retained file and are not reported here, and in any case a housekeeping promoter that varies across polarization states is not evidence that normalization succeeded.

### 2.4. Single Cell RNA-seq

#### 2.4.1. Datasets and Subsampling

Two published single-cell atlases were reanalysed. The liver dataset (GSE115469) comprises 8,444 cells and 19,563 genes from five healthy human donors, profiled by droplet-based 10x Genomics single-cell RNA-seq [28,29]. The blood dataset (GSE150728) comprises peripheral blood mononuclear cells from seven patients with severe COVID-19 and six healthy donors, profiled by Seq-Well [30,31]. Cell numbers per-donor group are as reported in each source publication; the deposited GEO series for the blood atlas carries sample records for six distinct patient identifiers.

To keep the blood object small enough to reanalyse repeatedly on a laptop, the full 44,721 cell atlas was reduced to a representative subsample of 8,006 cells (21,742 genes) by stratified proportional sampling within every condition by cell type stratum, with a floor of 15 cells per stratum and a fixed random seed (42). All 13 donors are retained and cell type composition is approximately preserved, except in strata that hit the 15 cell floor. The subsample reproduces the full atlas result: the proportion of MS4A4A-detected monocytes is 15.3% in COVID-19 and 4.1% in healthy donors in the subsample, against 15.4% and 5.0% in the full atlas. The sampling procedure and this comparison are recorded in the data provenance record supplied with Supplementary Software S1, which also contains the analysed count matrix and its per cell annotations. All single cell results reported here are from the subsample.

#### 2.4.2. Processing and Cell Type Labels

Both datasets were processed identically in Seurat 5.5.1 (SeuratObject 5.4.0) in R 4.6.1, with a fixed seed of 42: ‘NormalizeDatà, then ‘FindVariableFeatures’, ‘ScaleDatà, ‘RunPCÀ (30 principal components), ‘FindNeighbors’ (dimensions 1 to 20), ‘FindClusters’ (resolution 0.5), and ‘RunUMAP’ (dimensions 1 to 20) [32], the last of these embedding the cells by uniform manifold approximation and projection (UMAP) [33,34]. Quality control metrics, namely genes detected per-cell (’nFeature_RNÀ), total counts per-cell (’nCount_RNÀ), and percentage of mitochondrial transcripts (’percent.mt’), were computed and retained as cell-level metadata.

Cells are those retained in the published annotated datasets; no additional cell-level filtering was applied here, and genes were those detected in at least three cells in the source data.

Cell type assignments were taken from the published annotations accompanying each dataset rather than derived de novo. In the liver dataset, MacParland et al.’s two macrophage labels, "inflammatory macrophage" (822 cells) and "non inflammatory" or Kupffer macrophage (326 cells), were merged into a single Macrophage cell type (1,148 cells) for Figure 2a,b, and retained separately as M1-like and M2-like respectively for Figure 2c. Alpha beta and gamma delta T cells were merged into a single T cell label (1,786 cells).

**Figure 2.**
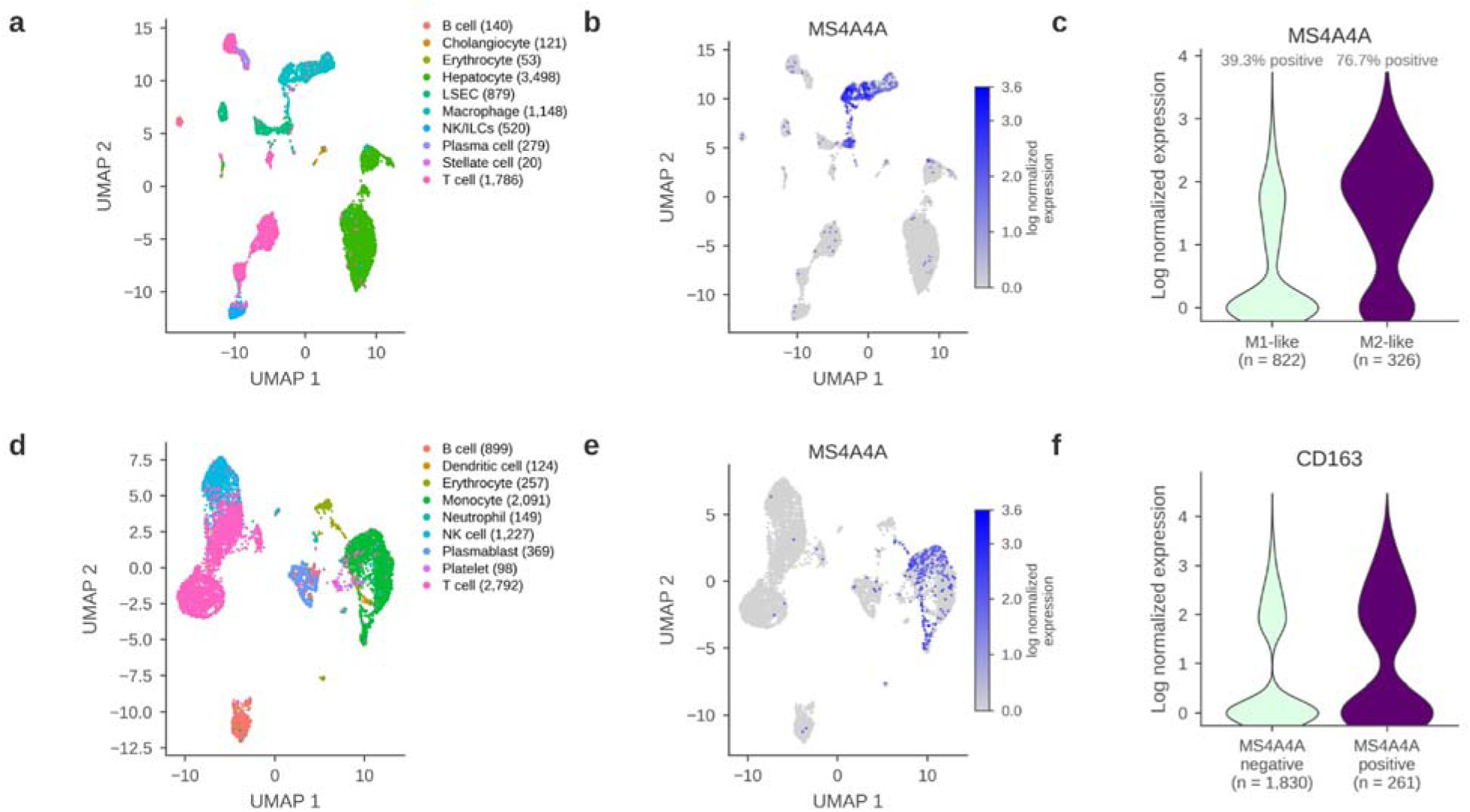
**MS4A4A marks the M2 side of the myeloid compartment in two independent human single-cell datasets**. **(a)** UMAP of 8,444 cells from healthy human liver (GEO GSE115469; five donors), coloured by the ten annotated populations, with group sizes given in the panel legend: macrophages (n = 1,148), liver sinusoidal endothelial cells (LSECs; 879), hepatocytes (3,498), cholangiocytes (121), stellate cells (20), T cells (1,786), NK/ILCs (520), B cells (140), plasma cells (279), and erythrocytes (53). Each point is one cell. **(b)** MS4A4A expression projected onto the same UMAP. Grey indicates no detected expression and darker blue indicates higher expression, on a scale of 0 to 3.6 log normalized units. Signal is concentrated over the macrophage cluster. **(c)** MS4A4A expression in liver macrophages, split by the atlas’s polarization annotation (M1-like = inflammatory macrophage, n = 822; M2-like = Kupffer cell macrophage, n = 326). Violin width shows the density of cells at each expression level.

#### 2.4.3. Embeddings, Projections, and Group Comparisons (Figure 2)

UMAP embeddings were coloured by cell type label (Figure 2a,d) or by *MS4A4A* expression (Figure 2b,e). Panels 2b and 2e use one explicit colour scale, 0 to 3.6 log normalized units, set from the larger of the two datasets’ maxima and passed to both panels, so the two are directly comparable.

For Figure 2c, macrophages were subset from the liver object and *MS4A4A* expression compared between the M1-like and M2-like groups by violin plot. A cell was defined as *MS4A4A* detected if at least one *MS4A4A* count was detected in the raw counts layer. The proportion of positive cells was computed per group, and *MS4A4A* expression was compared between groups by two-sided Wilcoxon rank sum test.

For Figure 2f, monocytes were subset from the blood object (n = 2,091) and split by MS4A4A detection status using the same definition of at least one count, giving 261 positive and 1,830 negative cells. Expression of CD163, which encodes a marker of M2 macrophages, was compared between MS4A4A-detected and MS4A4A-undetected monocytes by violin plot. Mean log normalized expression of three further markers, MERTK, C1QA, and the pro-inflammatory cytokine gene TNF, was compared between the same two groups and is reported as a fold difference in Table S8, together with a two-sided Wilcoxon rank sum test per marker. Detection frequencies for every group are given in Table S7.

### 2.5. Structure Prediction

#### 2.5.1. Sequences and Prediction

Canonical full length human sequences were retrieved from UniProt: MS4A4A (accession Q96JQ5; 239 residues) and MS4A6A (accession Q9H2W1; 248 residues) [35]. Both proteins are annotated with four transmembrane helices, MS4A4A at residues 65 to 85, 99 to 119, 138 to 158, and 180 to 200, and MS4A6A at residues 47 to 67, 85 to 105, 117 to 137, and 186 to 206.

Both structures were predicted with AlphaFold 3 via the AlphaFold Server [17]. Two predictions were made: the MS4A4A monomer, run on 29 July 2026 with random seed 58014515; and a two chain MS4A4A-MS4A6A complex, run on 30 July 2026 with random seed 1735480323, in which chain A is MS4A4A (residues 1 to 239) and chain B is MS4A6A (numbered 240 to 487 in the concatenated numbering used in Figure 3f). No model was taken from the AlphaFold Protein Structure Database, so the five predicted panels of Figure 3 all derive from the same model generation; panel 3b is an experimental structure. Predictions were made without a lipid bilayer, ligands, or post translational modifications. Five models were returned for each prediction and the top ranked model was used for all panels. Run parameters and confidence metrics are summarized in Table S12.

**Figure 3.**
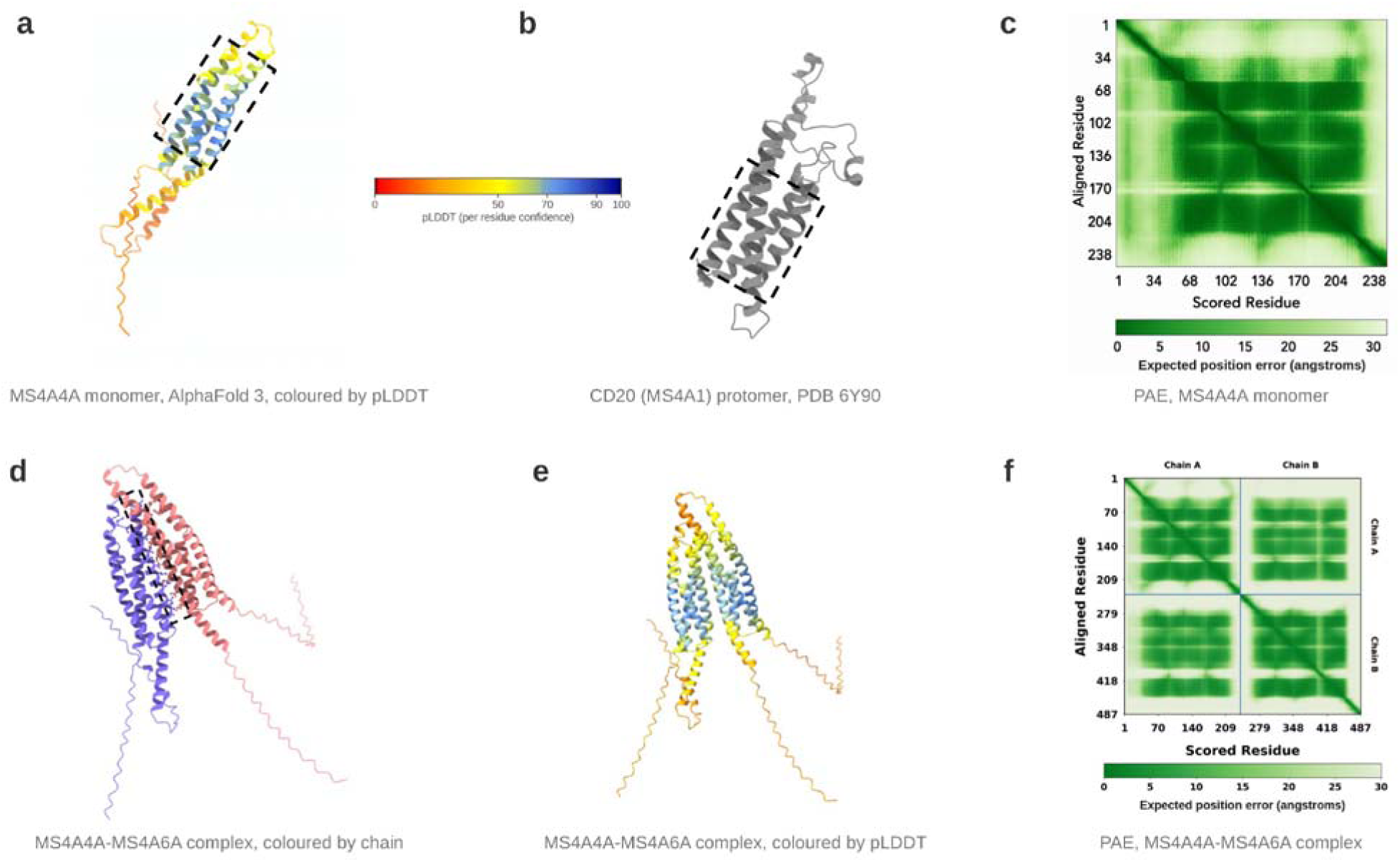
AlphaFold predicted structures of the MS4A4A monomer and the MS4A4A-MS4A6A complex. (a) Predicted structure of full length human MS4A4A (UniProt Q96JQ5; 239 residues) from AlphaFold 3, coloured by per-residue confidence (pLDDT, on the 0 to 100 scale keyed beside the panel; red is low, blue is high). The dashed box marks the four transmembrane helical core, corresponding to the helices annotated at residues 65 to 85, 99 to 119, 138 to 158, and 180 to 200. This core is the highest confidence region of the model (mean pLDDT 83.1, AlphaFold’s confident band); the extended N and C terminal segments outside it are predicted with low-confidence and bring the mean across the whole model to 62.4. (b) A single CD20 (MS4A1) protomer, extracted from the cryo-electron microscopy structure of the full length CD20 homodimer in complex with two rituximab Fab fragments (PDB 6Y90, 3.69 Å [15]) and shown in a comparable orientation. The dashed box marks its four transmembrane bundle. CD20 is the only member of the MS4A family for which an experimental structure has been determined. (**c**) Predicted aligned error (PAE) for the MS4A4A monomer, residues 1 to 239. Each cell reports the expected error in the position of the scored residue when the model is aligned on the aligned residue, on a 0 to 30 Å scale; darker indicates lower expected error. Error is lowest among residue pairs within the transmembrane core and highest for pairs involving either terminus. (**d**) Predicted MS4A4A-MS4A6A complex coloured by chain (MS4A4A blue, MS4A6A salmon). The dashed box marks the predicted inter-chain contact region, within which residues at the predicted interface are drawn as sticks; these are listed in Table S9, in each protein’s own residue numbering. Confidence in this arrangement is low: ipTM 0.59, pTM 0.62. Values of ipTM below approximately 0.6 are not conventionally taken to support a predicted interface, so the contact shown here is a hypothesis rather than a result. (**e**) The same predicted complex coloured by pLDDT, on the scale keyed in (**a**). The helical core of each chain is predicted with higher confidence than the extended terminal segments, reproducing the pattern seen for the monomer in (**a**). (**f**) PAE for the predicted complex, on the same 0 to 30 Å scale as (**c**). Chain A is MS4A4A (residues 1 to 239) and chain B is MS4A6A (UniProt Q9H2W1; 248 residues, numbered 240 to 487 here); the two chains are marked on the axes and separated by rules. The two on-diagonal blocks report within-chain residue pairs and the two off-diagonal blocks report between chain pairs; the off-diagonal blocks carry higher expected error than the on-diagonal blocks, modestly so when whole blocks are compared, which is the graphical form of the low ipTM in (**d**). All models shown are AlphaFold 3 predictions except (**b**), which is an experimental structure. Both predictions were run on the AlphaFold Server; run dates, seeds, and the confidence metrics recorded for each are in Table S12.

#### 2.5.2. Confidence Assessment

Model confidence was assessed using the predicted local distance difference test (pLDDT) [36] for per-residue confidence, and predicted aligned error (PAE) for confidence in the relative positioning of residue pairs. pLDDT is reported on a 0 to 100 scale, with values above 90 indicating very high-confidence, 70 to 90 confident, 50 to 70 low, and below 50 very low. PAE is reported in Å on a 0 to 30 scale in Figure 3c,f. Mean pLDDT was 62.44 across the MS4A4A monomer overall and 83.09 across its four annotated transmembrane helices. For the complex, the interface predicted template modelling score (ipTM) was 0.59 and the predicted template modelling score (pTM) was 0.62. ipTM is the diagnostic for whether a predicted inter-chain arrangement should be believed [20]. Published guidance places ipTM above 0.8 in the high quality band, 0.6 to 0.8 in a grey zone, and below 0.6 among likely failed predictions, with benchmarking placing the discriminative cut off between 0.65 and 0.75 [37]; 0.59 falls below the permissive end of that range, and we interpret the MS4A4A-MS4A6A complex accordingly throughout.

#### 2.5.3. Comparison with an Experimental MS4A Structure

The predicted MS4A4A monomer was compared with CD20 (MS4A1), the only MS4A family protein for which an experimental structure has been determined. A single CD20 protomer was extracted from PDB 6Y90, the cryo-electron microscopy structure of the full length CD20 homodimer bound to two rituximab Fab fragments at 3.69 Å resolution, and oriented for visual comparison [15]. The two models were not formally superposed and no C alpha root mean square deviation is reported, so the correspondence between the predicted MS4A4A fold and the experimental CD20 fold is stated qualitatively rather than quantitatively.

#### 2.5.4. Visualization and Interface Definition

Predicted models were visualized in UCSF ChimeraX 1.12 [38]. The residues drawn as sticks inside the dashed contact region in Figure 3d, and the criterion used to select them, are given in Table S9; residues of chain B are listed there in MS4A6A’s own residue numbering, 1 to 248, rather than in the concatenated numbering used on the axes of Figure 3f. Because the complex is a computational prediction with ipTM 0.59, these are reported as candidate contacts rather than validated interaction sites.

### 2.6. Statistical Analysis

Differential expression in bulk RNA-seq was assessed by Wald test in DESeq2 with Benjamini and Hochberg correction across genes; significance was defined as adjusted *P* < 0.05 with absolute log_2_ fold change > 1. Single cell expression between two groups was compared by two-sided Wilcoxon rank sum test across individual cells. Such a test treats cells from the same donor as independent observations, which inflates significance and can return large numbers of differentially expressed genes in the absence of any biological difference between groups [39]; the very small *P* values it returns therefore reflect the number of cells as much as the size of the effect, and the effect size and the difference in detection fraction are the more informative quantities. No donor-level aggregation or mixed model was applied, either of which would be the stricter approach [39,40]. The disease stratified monocyte percentages were not tested at all, for the same reason. ATAC-seq accessibility is reported descriptively as fold over genome mean per-donor; with three donors per state, no significance test was applied. Structural confidence measures (pLDDT, PAE, pTM, ipTM) are model diagnostics, not statistical tests. No adjustment for multiple comparisons was made across figures.

### 2.7. Software and Use of Generative Artificial Intelligence

R 4.6.1; DESeq2 1.52.0; EnhancedVolcano 1.30.0; ggrepel 0.9.8; Seurat 5.5.1; SeuratObject 5.4.0; ggplot2 4.0.3; dplyr 1.2.1; tidyverse 2.0.0 (including readr 2.2.0, tidyr 1.3.2, purrr 1.2.2, stringr 1.6.0, forcats 1.0.1, lubridate 1.9.5); patchwork 1.3.2; Matrix 1.7-6; Python 3 with pydeseq2 0.5.2 and the standard library modules ‘struct’ and ‘zlib’; AlphaFold 3 (AlphaFold Server); UCSF ChimeraX 1.12. Random seeds were fixed at 42 for all stochastic steps in the R and Python analyses; the two AlphaFold seeds are given in Table S12. Per panel ‘sessionInfo()’ output is provided in Supplementary Software S1 for the Figure 1 panels; R 4.6.1 was used throughout, on two machines (Figure 1a,b and all of Figure 2 on macOS, Figure 1c,d on Windows 11). No session record was captured for the Figure 2 panels, so the Seurat and SeuratObject versions listed in Table S13 were read from the loaded namespace list of the Figure 1a record rather than from a session record of their own. Versions are summarized in Table S13.

Generative artificial intelligence was used in the preparation of this work, and the disclosure required by the journal is given in full in the Acknowledgments. It was used to assist with drafting and editing of manuscript text, with code for figure generation, and with reference checking. It was not used to generate, alter, or select any data, and every numerical value reported in this paper was computed by the analysis scripts provided in Supplementary Software S1. All output was reviewed and verified by the authors, who take full responsibility for the content.

## 3. Results

### 3.1. *MS4A4A* Is Upregulated in M2 Macrophages, and Its Promoter Is More Accessible

To ask whether *MS4A4A* differs between the two best characterized macrophage states, we reanalysed bulk RNA-seq of primary human monocyte-derived macrophages polarized to M1 (IFN-γ + LPS) or M2 (IL-4) from three donors (GSE162698) [22]. Across 12,900 genes, 1,534 were higher in M2 and 1,692 were higher in M1 (adjusted *P* < 0.05, absolute log_2_ fold change > 1), and the most strongly M2 enriched transcripts included the canonical IL-4 response genes *ALOX15*, *CCL13*, *CD1C*, and *CLEC10A* (Table S2; the volcano plot itself labels only *MS4A4A*). *MS4A4A* fell on the M2 side of the volcano plot and passed both thresholds, with a log_2_ fold change of +2.68 and an adjusted *P* value of 0.0013, approximately a six fold increase in M2 relative to M1 (Figure 1a). *MS4A4A* is not among the most extreme genes on either axis: it ranks 2,427th of 12,900 genes by adjusted *P* value, sitting above the significance cut off but well below the top ranked transcripts.

Notably, *MS4A6A*, the gene encoding the proposed structural partner examined in Figure 3, was co induced in the same contrast and more strongly than *MS4A4A*, with a log_2_ fold change of +4.03 and an adjusted *P* value of 1.2 × 10^-4^ (Table S2). The two genes therefore move together across this polarization axis.

Because a genome-wide test can be driven by one sample, we next examined *MS4A4A* donor by donor (Figure 1b; Table S3). Expression was higher in M2 than in M1 in all three donors, rising from 2,380 to 8,769 pseudocounts in donor 1, from 75 to 2,962 in donor 2, and from 329 to 2,061 in donor 3. M1 expression varied more than thirty fold between donors, and the two groups overlap, since the highest M1 value (2,380) exceeds the lowest M2 value (2,061), so the within donor pairing, rather than the separation of the two distributions, is what carries the signal. The analysis reflects this: the donor-blocked model gave a log_2_ fold change of +2.68 (adjusted *P* = 0.0013), whereas the same contrast without donor blocking gave +2.33 (adjusted *P* = 0.027) (Table S4).

We then asked whether this difference in transcript level is mirrored in chromatin. ATAC-seq from an independent set of primary human monocytes and monocyte-derived macrophages (GSE261697; monocytes and M0, M1, and M2 (IL-4) macrophages from three donors) showed a discrete accessibility peak at the *MS4A4A* promoter in every state, and M2 showed the highest mean signal and the lowest between-donor variability among the three macrophage states across the three donors analysed (Figure 1c). At the promoter apex the donor mean signal reached 29.7 fold over genome mean in M2, against 17.9 fold in M1, 19.1 fold in M0, and 12.1 fold in monocytes. Two further peaks inside the gene body, at approximately chr11:60,049,900 and chr11:60,053,000, were also highest in M2 (14.5 and 12.2 fold over genome mean, versus 4.4 and 0.7 fold in monocytes), whereas a strong peak immediately downstream of the gene (chr11:60,078,200) was present in all three macrophage states (29.6, 16.6, and 21.9 fold in M2, M1, and M0) and, at lower level, in monocytes (6.0 fold) (Table S5).

Quantifying the promoter window for each donor separately made the pattern explicit (Figure 1d; Table S6). Monocytes were uniformly low (2.73, 2.28, and 1.96 fold over genome mean in donors A, B, and C; mean 2.33, SD 0.39). Differentiation to M0 and polarization to M1 raised the mean accessibility, to 4.66 and 4.28 respectively, but did so inconsistently: in M0, donor B reached 10.59 while donors A and C remained at 2.67 and 0.73; in M1, donor A reached 7.78 while donors B and C remained at 2.92 and 2.13. M2 gave both the highest mean (9.17, approximately four fold above monocytes) and the narrowest spread across donors of any macrophage state (7.29 to 10.57; SD 1.69, versus 5.23 for M0 and 3.06 for M1; monocytes, which are uniformly low, are tighter still at SD 0.39). M2 promoter accessibility exceeded M1 in every donor, by 2.79, 6.73, and 5.15 fold over genome mean units. Across states, M2 was the state with the highest mean signal in donors A and C; in donor B, M0 was marginally higher (10.59 versus 9.65).

Across two independent donor cohorts and two different assays, therefore, MS4A4A transcript level and MS4A4A promoter accessibility were both higher in M2 than in M1. Promoter mean accessibility signal was highest in M2 of the four states assayed, in these three donors; transcript level was not, because MS4A4A is also high in M0 and in TAM-like macrophages (Table S3) and low specifically in M1.

### 3.2. *MS4A4A* Is Macrophage Enriched in Human Liver and Is Expressed by More M2-Like than M1-Like Macrophages

To locate MS4A4A among the cell types of a human tissue, we analysed a published single-cell RNA-seq atlas of healthy human liver (GSE115469; 8,444 cells from five donors) [28]. The published annotation carries twelve labels which, with the two macrophage labels and the two T cell labels merged (see Section 2.4.2), give ten populations: macrophages (n = 1,148), liver sinusoidal endothelial cells (LSECs; 879), hepatocytes (3,498), cholangiocytes (121), stellate cells (20), T cells (1,786), natural killer cells and innate lymphoid cells (NK/ILCs; 520), B cells (140), plasma cells (279), and erythrocytes (53). These separated cleanly on the UMAP (Figure 2a). Projecting MS4A4A onto the same coordinates placed almost all of the signal over the macrophage cluster (Figure 2b). Scattered positive cells appeared in other populations, most visibly among LSECs, where 11.9% of cells (105 of 879) carried at least one MS4A4A transcript, but macrophages were the only population in which MS4A4A-detected cells formed a broad, continuous distribution rather than isolated points. MS4A4A is therefore macrophage-enriched in this tissue rather than exclusive to macrophages.

MS4A4A was detected in 76.7% of M2-like and 39.3% of M1-like macrophages; expression differed between the two groups by two-sided Wilcoxon rank sum test (P = 4.5 × 10-39, computed across cells, see Section 2.6). **(d)** UMAP of 8,006 blood cells from seven patients with severe COVID-19 and six healthy donors (GEO GSE150728; representative subsample, see Section 2.4.1), coloured by the nine annotated populations: T cells (n = 2,792), monocytes (2,091), NK cells (1,227), B cells (899), plasmablasts (369), erythrocytes (257), neutrophils (149), dendritic cells (124), and platelets (98). **(e)** MS4A4A expression projected onto the same UMAP, on the identical colour scale used in (b). Signal is concentrated in the monocyte cluster; a few positive cells appear elsewhere. **(f)** Expression of CD163, which encodes a marker of M2 macrophages, in monocytes (n = 2,091) split by MS4A4A detection status: 1,830 negative and 261 positive. Monocytes were called MS4A4A detected if at least one MS4A4A transcript was detected. Violin width shows the density of cells at each expression level; the two fill shades in (c) and (f) separate the groups visually and carry no other meaning. Expression values in (b,c,e,f) are log normalized counts. Because single-cell dropout causes truly expressing cells to read as zero, the percentages of positive cells are underestimates. Cell type and polarization labels are the published annotations of the source atlases, not calls derived here.

Within the macrophage compartment, we compared the cells carrying the atlas’s M1-like (inflammatory macrophage; n = 822) and M2-like (Kupffer cell macrophage; n = 326) annotations (Figure 2c). *MS4A4A* was detected in 76.7% of M2-like macrophages (250 of 326 cells) and in 39.3% of M1-like macrophages (323 of 822 cells), and *MS4A4A* expression differed between the two groups by two-sided Wilcoxon rank sum test (*P* = 4.5 × 10^-39^; Table S7). The test compares expression values across cells rather than the two proportions, and treats cells as independent observations (see Section 2.6). The two violins reach a similar height: among the cells that are positive, the per-cell expression level is comparable in the two states, at mean log normalized values of 1.95 in M2-like and 1.67 in M1-like macrophages. What differs is the mass of cells at zero, which is far larger on the M1-like side. The difference between the two states is therefore driven principally by how many macrophages express *MS4A4A* rather than by how much each expressing macrophage makes.

### 3.3. In Blood, *MS4A4A* Is Monocyte Enriched, and *MS4A4A* Positive Monocytes Carry More M2 Associated Markers

To ask whether the same association holds outside solid tissue, we analysed a published single-cell atlas of peripheral blood from seven patients with severe COVID-19 and six healthy donors (GSE150728), subsampled to 8,006 representative cells (see Section 2.4.1) [30]. The published annotation resolved nine populations: T cells (n = 2,792), monocytes (2,091), NK cells (1,227), B cells (899), plasmablasts (369), erythrocytes (257), neutrophils (149), dendritic cells (124), and platelets (98) (Figure 2d). *MS4A4A* expression was again concentrated in a single population, the monocytes, where it reached values of 1 to 3 on the log normalized scale, while T cells, NK cells, and B cells were essentially negative (Figure 2e). A few positive cells appeared among dendritic cells and, more sparsely, in the remaining populations.

Because MS4A4A marked only part of the monocyte pool, with 261 of 2,091 monocytes (12.5%) carrying at least one transcript, we divided monocytes into MS4A4A-detected (n = 261) and MS4A4A-undetected (n = 1,830) cells and asked how the two groups differed. CD163, which encodes a haemoglobin scavenger receptor and canonical marker of alternatively activated macrophages [41], was higher in MS4A4A-detected monocytes, by 1.41 fold on average (Figure 2f; Table S8). CD163 was zero-heavy in both groups, with a large fraction of cells at zero and a second lobe near 2 on the log normalized scale; the MS4A4A-detected monocytes carried proportionally more of their cells in that positive lobe (50.6% versus 36.9% of cells above zero). Two further markers of the resolving, efferocytic state ran in the same direction [42], MERTK 2.53 fold and C1QA 3.92 fold higher in MS4A4A-detected monocytes, whereas the pro-inflammatory cytokine gene TNF was 0.47 fold, that is lower, in the same cells. Cell level Wilcoxon rank sum tests support the three markers that are higher (CD163 P = 1.1 × 10-5, MERTK P = 3.1 × 10-5, C1QA P = 2.0 × 10-4) but not the lower TNF value (P = 0.09), so the TNF difference should be read as a trend rather than as a result. These tests carry the same pseudoreplication caveat as the liver comparison (see Section 2.6).

The blood atlas also carries a disease label. Among the 2,091 monocytes analysed here (1,560 from patients with COVID-19, 531 from healthy donors), the proportion positive for MS4A4A was 15.3% in COVID-19 and 4.1% in healthy donors; the corresponding values in the full 44,721 cell atlas are 15.4% and 5.0% (see Section 2.4.1). We report this observation because it locates the MS4A4A-detected monocyte population in a disease setting, but it is a between group comparison in pooled cross sectional data and no formal test of the difference was performed here. *MS4A4A* expression is thus enriched in macrophages in liver and in monocytes in blood, and within each of those compartments it is associated with the M2, anti-inflammatory end of the phenotypic range.

### 3.4. AlphaFold Predicts a Four Transmembrane Fold for MS4A4A, and an MS4A4A-MS4A6A Interface of Low Confidence

Prediction of full length MS4A4A (UniProt Q96JQ5, 239 residues) produced a compact bundle of four transmembrane alpha helices matching the four-helical segments annotated for this protein at residues 65 to 85, 99 to 119, 138 to 158, and 180 to 200 (Figure 3a) [35]. These helices carried the highest per-residue confidence in the model, with a mean pLDDT of 83.1 across the four annotated helices, inside AlphaFold’s confident band (70 to 90) though below its very high band (above 90). The N terminal segment preceding the first helix, the three interhelical loops, and the C terminal segment following the fourth helix were all predicted with much lower confidence, and the two termini adopt extended conformations; because the residues outside the four helices account for 155 of the 239, they pull the mean pLDDT for the model as a whole down to 62.4, which falls in the low band; their own implied mean is approximately 51. The confidence in this model is therefore concentrated almost entirely in the four-helix core, which is a third of the sequence. Placed in a comparable orientation, a single protomer of CD20, also called MS4A1, and the only member of the MS4A family with an experimentally determined structure, showed the same four transmembrane organization (Figure 3b), providing an external reference for the predicted fold [15].

The PAE map for the monomer was consistent with this reading (Figure 3c). Expected positional error was lowest for residue pairs lying within the four-helix core, indicating that the model places these helices confidently with respect to one another, and was highest for pairs involving either terminus, whose positions the model does not resolve.

A model of the MS4A4A-MS4A6A complex was also predicted, in which the transmembrane regions of the two chains pack against one another across a contact surface (Figure 3d).

Coloured by confidence, both chains reproduced the pattern seen for the monomer: well predicted helical cores and poorly predicted terminal extensions (Figure 3e). The complex PAE map, however, separates the two kinds of information in the model (Figure 3f). The on-diagonal blocks, reporting residue pairs within MS4A4A (residues 1 to 239) or within MS4A6A (UniProt Q9H2W1, 248 residues, numbered 240 to 487 here), show low expected error. The off-diagonal blocks, reporting pairs that span the two chains, carry higher expected error than the on-diagonal blocks, and are worst for pairs involving the terminal segments. The difference between whole blocks is modest on the 0 to 30 Å scale, and is clearest for pairs between the two helical cores; ipTM is the quantitative statement of the same thing, and it is what we rely on. The prediction is therefore confident about the fold of each protein individually and considerably less confident about how the two are positioned relative to one another. The interface score for the model agrees: ipTM was 0.59. The overall pTM was 0.62, itself below the value usually taken to indicate a confidently placed overall fold, so the per chain confidence rests on pLDDT and on the on-diagonal PAE blocks rather than on pTM. Because ipTM values below approximately 0.6 are generally not treated as evidence of a correctly predicted interface [37], this model does not support the proposed MS4A4A-MS4A6A arrangement; it is a hypothesis of the right kind, generated at a confidence level too low to be relied upon. The residues drawn at the predicted interface in Figure 3d are accordingly reported as candidate contacts rather than as validated interaction sites, and are listed in Table S9. They are informative about the shape of the proposed arrangement even though its geometry is not supported: in both chains the contacts fall in the first and fourth transmembrane helices and in the extracellular loops flanking them, and the second and third helices contribute none. The model therefore packs the two four-helix bundles edge to edge through one face of each, rather than burying all four helices, and the contact is weighted towards the extracellular side of the membrane.

## 4. Discussion

We integrated bulk transcriptomic, chromatin accessibility, single cell transcriptomic, and structure prediction analyses to describe MS4A4A across four levels of biological organization: when it is expressed, whether its regulatory region is open, which individual cells express it, and what shape the encoded protein may take. Rather than relying on one kind of evidence, we asked whether independent assays applied to independent cohorts converge. They do: the three transcriptomic analyses and the chromatin analysis agree in associating MS4A4A with the M2 side of macrophage activation; the structural analysis addresses fold, not M2 association. Figure 4 summarizes that convergence and, equally deliberately, marks the two places where it stops.

**Figure 4.**
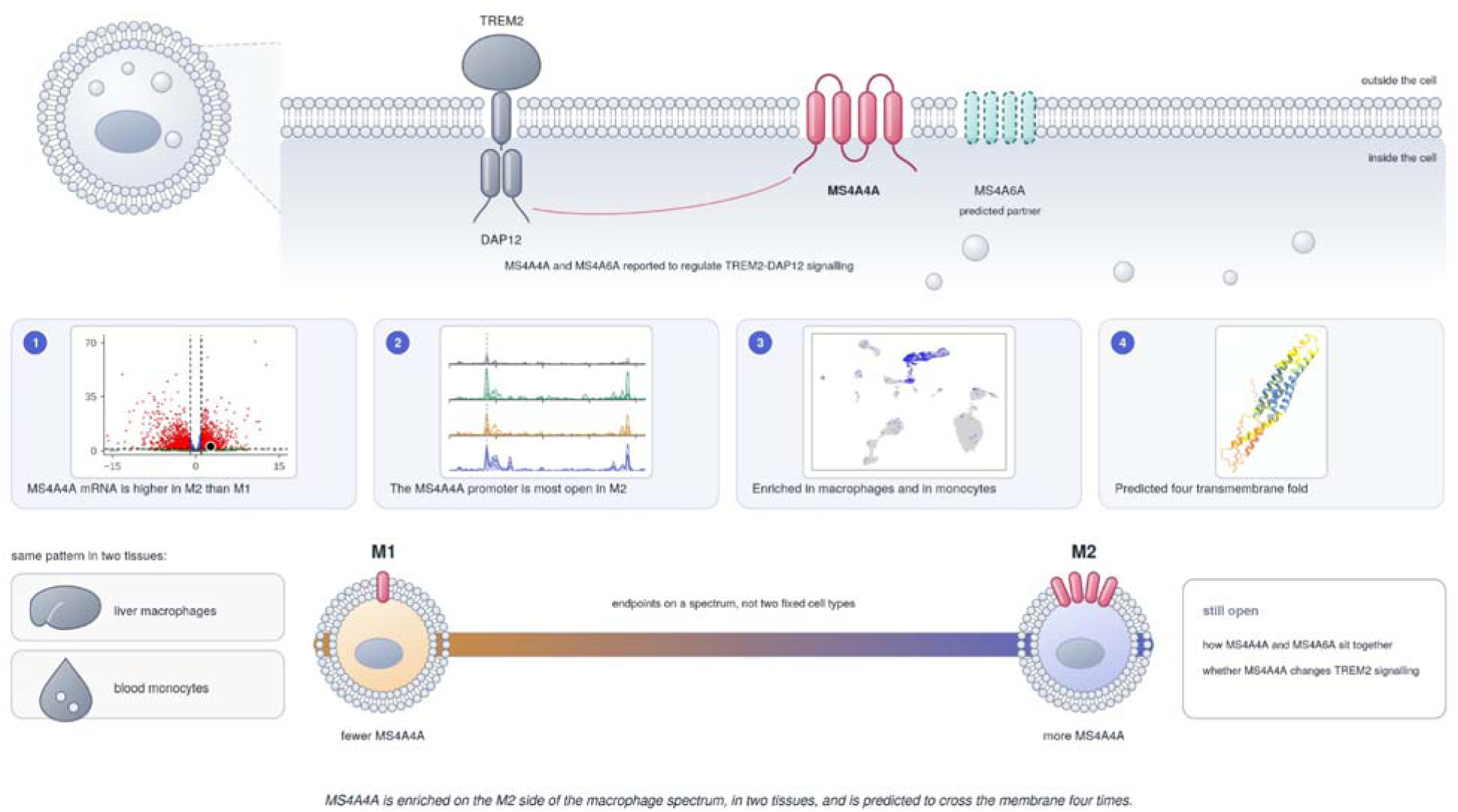
Integrated summary of MS4A4A expression, chromatin accessibility, cellular localization, and predicted membrane topology. Top: Schematic of MS4A4A in the macrophage plasma membrane. MS4A4A is shown as a four-pass transmembrane protein, consistent with its annotated topology and the AlphaFold 3 prediction in Figure 3a. MS4A6A is shown as a predicted partner with a dashed outline because the AlphaFold-predicted MS4A4A–MS4A6A complex had low interface confidence (ipTM = 0.59) and therefore does not establish their relative arrangement or interaction. The connection to TREM2–DAP12 represents a previously reported regulatory relationship [13] and was not tested in this study. **Middle:** Four findings summarized from Figures 1–3: (1) MS4A4A mRNA is higher in M2 (IL-4) than in M1 (IFN-γ + LPS) macrophages; (2) MS4A4A promoter accessibility shows the highest mean signal in M2 among the four states examined in three donors; (3) MS4A4A is enriched in liver macrophages and blood monocytes and is associated with M2-like myeloid states; and (4) AlphaFold predicts a four-transmembrane-helix core for MS4A4A. **Bottom:** Integrated model placing MS4A4A toward the M2 side of a continuous macrophage activation spectrum rather than defining M1 and M2 as discrete cell types. The schematic representation of fewer MS4A4A molecules on the M1 side and more on the M2 side is a simplified visual summary; in the liver single-cell dataset, the observed difference primarily reflected a higher proportion of MS4A4A-detected cells in the M2-like than in the M1-like macrophage population (76.7% versus 39.3%), whereas expression among positive cells was comparable between the two groups. The liver and blood symbols indicate that the association was observed in liver macrophages and blood monocytes. The MS4A4A–MS4A6A arrangement and any effect of MS4A4A on TREM2 signalling remain unresolved. This schematic summarizes results from Figures 1–3 and cited prior work, contains no new data, and should not be interpreted as evidence of causality.

### 4.1. Transcript Level and Chromatin Accessibility Agree

Bulk RNA-seq of primary human monocyte-derived macrophages showed *MS4A4A* significantly increased in M2 (IL-4) relative to M1 (IFN-γ + LPS), by approximately six fold. What makes this result interpretable is not the fold change but the donor structure behind it: expression rose from M1 to M2 in every donor despite more than thirty fold variation in M1 expression between donors, and the M1 and M2 ranges overlap, so the effect is carried by within donor pairing rather than by separation of the two distributions. The donor-blocked estimate was sharper than the unblocked one, as expected when donor to donor variance is removed from the residual, though the unblocked run also used a different implementation (see Section 2.2.4), so the two differ in more than the design. Either way, the pattern underscores the value of paired designs in primary human macrophage work [1,3].

ATAC-seq from an independent donor cohort found a discrete *MS4A4A* promoter peak in every state but the highest and most reproducible signal in M2, with the narrowest inter donor spread of any macrophage state and higher accessibility in M2 than M1 in every donor. Two intragenic peaks were also M2 biased, suggesting regulation is not confined to a single promoter element. This is the pattern expected if chromatin at the locus opens in step with IL-4 signalling through STAT6 [43], and with the histone H3K27 demethylation that permits IRF4 expression in M2 macrophages [44], although these data cannot demonstrate either mechanism, and neither study examined this locus.

It is worth being explicit about what these two assays do and do not establish jointly. ATAC-seq reports regulatory potential, not active transcription; RNA-seq reports transcript abundance, not its regulatory cause. Their agreement is therefore corroboration by independent measurement, not a demonstration of a mechanistic chain, and because they were performed on different donor cohorts, they are consistent with one another rather than matched. A single cell or bulk multi-omic design in matched cells would be required to link the two directly.

One asymmetry in the bulk data deserves comment. *MS4A4A* is also high in M0 macrophages and in tumour conditioned (TAM-like) macrophages, and is low specifically in M1. The M2 versus M1 gap is therefore driven substantially by suppression in M1 rather than by induction in M2 alone. This is consistent with *MS4A4A* behaving as part of a default myeloid maturation programme that pro-inflammatory signalling switches off, rather than as an IL-4 specific induced gene, and it is a distinction that a two state comparison cannot resolve.

We also note that *MS4A6A*, the gene encoding the proposed structural partner examined in Figure 3, was co induced with *MS4A4A* in the same contrast and to a greater degree. That the two genes are coordinately regulated across this axis makes the structural question in Figure 3 better motivated. Two caveats belong with that statement, though. These are bulk measurements of a polarized population, so they show that both transcripts rise in the same condition, not that both are expressed in the same individual cells; *MS4A6A* was not examined in either single-cell dataset. And they are transcripts, not proteins. Co regulation of this kind is a reason to ask the structural question; it is not evidence that the two proteins interact.

### 4.2. Single Cell Data Localize the Signal and Change Its Interpretation

Single cell RNA-seq of human liver placed *MS4A4A* predominantly in macrophages, with scattered signal in LSECs and essentially none in hepatocytes, T cells, or B cells. Within the macrophage compartment, *MS4A4A* was detected in 76.7% of M2-like versus 39.3% of M1-like cells. The important detail is that per-cell expression among positive cells was comparable between the two states; the difference lay in the fraction of cells above zero. *MS4A4A* therefore marks a larger M2 leaning subset rather than being uniformly dialled up in every M2 cell, a distinction that bulk analysis cannot make. It is consistent with switch-like rather than graded regulation, though it does not demonstrate it: in unique molecular identifier based droplet data the observed zeros are largely explained by sampling at limited depth rather than by a separate zero inflation process [45], so a detection difference cannot by itself distinguish a bimodal population from a graded one measured shallowly. If the switch-like reading is correct it would be a useful property in a marker, because such genes discriminate populations better than smoothly varying ones.

Blood single-cell data reproduced the myeloid enrichment on a different platform, in a different tissue, and in a patient cohort: MS4A4A was concentrated in monocytes, and MS4A4A-detected monocytes carried more CD163, MERTK, and C1QA than their negative counterparts, and less TNF, although the TNF difference alone did not reach significance. Recovering the same association in liver macrophages and blood monocytes argues that the link between MS4A4A and an M2-associated programme is a general property of human myeloid cells rather than a feature of one dataset, while acknowledging that M1/M2 categories are simplified descriptions of a broader activation continuum [1].

Two features of these data constrain how far the percentages can be pushed. Single cell dropout causes genuinely expressing cells to read as zero, so every positive fraction reported here is an underestimate; because the liver M1-like and M2-like difference is itself a difference in detection frequency, this matters more than usual. The accompanying Wilcoxon *P* values are computed across cells and so treat cells from one donor as independent; they should be read as descriptions of the observed cell distributions rather than as inferences about donors. That the liver difference is nonetheless large and highly significant supports a conservative reading. Conversely, systematic comparisons of single-cell methods place Seq-Well below droplet-based 10x Genomics in transcripts and genes detected per-cell [46], so absolute detection rates in the blood atlas are not comparable to those in the liver atlas, and we do not compare them.

### 4.3. The Structural Prediction Is Confident about the Fold and Not about the Interface

AlphaFold 3 predicted MS4A4A as a compact four-pass transmembrane protein, confidently modelled in the helical core (mean pLDDT 83.1) and poorly modelled at the termini. The whole model mean of 62.4 is worth stating alongside it, because reporting only the core figure would flatter the model badly: 155 of the 239 residues, roughly two thirds of the protein, lie outside the four helices in long extended segments, and their implied mean pLDDT is about 51, at the edge of AlphaFold’s very low band. This model resolves a confidently placed four-helix bundle; the 64 residue N terminus and the 39 residue C terminus that hang off it, and the loops between the helices, it does not resolve at all. The predicted core agrees with the topology annotated for the protein and is visually consistent with the experimentally determined fold of CD20/MS4A1, which supports the model at the level of the individual protein. That comparison was made by orienting the two models similarly rather than by formal superposition, so it should be read as qualitative. Agreement between pLDDT, which reports local confidence, and the monomer PAE map, which reports confidence in relative positioning, strengthens the reading of the core.

The predicted MS4A4A-MS4A6A complex must be read differently. Both chains individually reproduced the monomer’s confidence pattern, but inter-chain expected error was elevated relative to within-chain error, and highest for every pair involving a terminal segment. The elevation is modest when whole blocks are compared, which is why the argument rests on ipTM rather than on the appearance of the map. The decisive measure is ipTM, which was 0.59, below the threshold at which a predicted interface is conventionally treated as supported [37].

We take the honest reading: the model is informative about two folds and uninformative about their arrangement. The residues shown at the interface are candidate contacts requiring experimental test, not established binding sites. We report the low score rather than omitting it, because a predicted complex presented without an interface score is indistinguishable to a reader from one that was actually supported. This illustrates a general principle worth stating plainly: high within-chain pLDDT does not license belief in a predicted docking geometry, and for multi chain predictions the off-diagonal PAE block, together with ipTM, is the diagnostic that separates confidence in the parts from confidence in the assembly [20,37].

One piece of independent structural context is worth stating precisely, because it is easy to overstate: CD20/MS4A1 is itself a homodimer in the experimental cryo-electron microscopy structure, associating through its transmembrane helices [15]. That establishes that at least one MS4A protein can associate through its membrane embedded region, which makes MS4A heteromerization structurally conceivable. It is not evidence for this particular pair, since CD20 dimerizes with itself, and homodimerization of one paralogue does not predict heterodimerization of two others.

The reported relationship between MS4A4A, MS4A6A, and TREM2-DAP12 signalling is the most interesting downstream implication of this work and also the furthest from it. Genetic evidence links the *MS4A* locus to soluble TREM2 and to Alzheimer’s disease risk [12], and MS4A4A and MS4A6A have been shown to cooperate in negatively regulating TREM2 via DAP12 [13]. We measured no binding and no signalling; any mechanistic connection between our observations and that pathway remains speculative.

### 4.4. Limitations

#### 4.4.1. Data and Analytical Limitations

All data are public and were generated by other groups under different protocols and on different platforms, which introduces heterogeneity we cannot control. The bulk RNA-seq count matrix is derived from published gene-level TPM rather than raw sequencing counts, converted to integer pseudocounts for compatibility with DESeq2’s negative binomial model. That model assumes raw counts, so the exact *P* values reported for Figure 1a should be read as approximate; the direction and approximate magnitude of the *MS4A4A* effect are robust to this. A publication grade reanalysis would use true gene-level counts, via recount3 [47] or ARCHS4 [48] or realignment of the underlying Sequence Read Archive submission, or a method designed for continuous normalized values such as limma-trend [49,50]. The ATAC-seq values are normalized coverage, not called peaks, and the RNA-seq and ATAC-seq cohorts comprise different donors. Only three donors were studied per bulk assay, and bulk ATAC-seq cannot distinguish whether greater M2 accessibility reflects modest opening in most cells or extreme opening in a subset; single-cell ATAC-seq would resolve this and would allow direct RNA and chromatin correlation in the same cells. Coverage is also uneven between samples, as the quality control paragraph in Section 2.3.4 sets out: in the lowest sample (donor C, M0, promoter mean 0.73) 16 of the 26 bins in the promoter window carry exactly zero coverage, including the bin at the promoter apex, and 54.5% of all 420 bins across the locus are zero; the monocyte sample from donor A is similar. Low values in those samples may therefore reflect sequencing depth as much as closed chromatin, and because donor C’s M0 value is what makes the M0 group mean low and its standard deviation large, this qualifies the comparison of M2’s reproducibility against M0’s. It does not weaken the comparison against M1, whose three samples are as completely covered as M2’s.

#### 4.4.2. Annotation Limitations

The M1-like and M2-like labels used in Figure 2c are the liver atlas authors’ own annotations, "inflammatory macrophage" and Kupffer cell macrophage, reinterpreted onto the M1/M2 axis. They are not an independent polarization call derived from these data. Similarly, all cell type labels in both single-cell datasets were taken from the published annotations rather than assigned de novo. The blood dataset pools cells from patients with COVID-19 and healthy donors for the MS4A4A-detected versus negative comparison, so that comparison describes MS4A4A-detected monocytes in general and not a disease specific population; the disease stratified percentages are reported descriptively and untested. Of the four markers examined in MS4A4A-detected monocytes, only CD163 is shown as a panel; MERTK, C1QA, and TNF are reported as mean fold differences without accompanying plots.

#### 4.4.3. Structural Limitations

No experimental structure of MS4A4A exists, so every structural statement about MS4A4A in this paper is a prediction. Both models were predicted in isolation, without a lipid bilayer, ligands, or post translational modifications, so helix packing and loop conformations may differ in a membrane environment, and the long low-confidence terminal segments should not be interpreted as real conformations. Inter chain predicted aligned error is elevated across the MS4A4A-MS4A6A interface and ipTM is 0.59, below the conventional threshold, so the relative orientation of the two chains is not established by this model; the modelled interface should be treated as a target for experiment, not as a finding. The whole model mean pLDDT of 62.4 also means that most of the sequence outside the four helices is not modelled to a usable standard. AlphaFold does not model membrane dynamics or lipid effects; predictions with full MS4A family multiple sequence alignments, molecular dynamics in explicit bilayers, orthogonal predictors such as RoseTTAFold or ESMFold, and experimental distance constraints from cross-linking or hydrogen deuterium exchange would each add information [51,52].

#### 4.4.4. Functional Limitations

Every conclusion here is correlative. Whether MS4A4A drives M2-like polarization, helps maintain the state once established, or passively reflects it is not addressed by any of these data. Taken together, the limitations of this study are the following. It reanalyses public datasets generated in different biological and technical contexts and by different groups. Donor numbers are modest throughout: three donors per bulk assay, five liver donors and thirteen blood donors. Single-cell macrophage-state interpretations were based on source annotations and marker patterns rather than experimentally induced or functionally validated here. Droplet and Seq-Well single-cell measurements are affected by incomplete transcript detection, so every detected fraction is an underestimate. The bulk differential expression was computed from published TPM converted to pseudocounts rather than from count-level input, so its P values are approximate rather than confirmatory. The single-cell comparisons were tested across cells rather than across donors and are therefore descriptive. The ATAC-seq analysis is descriptive and covers three donors with uneven coverage. Finally, the structural analyses are predictions; the MS4A4A-MS4A6A model had limited interface confidence (ipTM 0.59) and does not establish a biochemical interaction. These results therefore motivate donor-matched validation, perturbation studies, protein-level localization and direct interaction assays.

### 4.5. Future Directions

The most direct next steps follow from the two questions this work leaves open. The first is whether MS4A4A and MS4A6A associate at all, and the order of experiments matters here because ipTM 0.59 makes the modelled geometry a weak prior. Association should be established first, by co immunoprecipitation, proximity ligation, or Forster resonance energy transfer. Geometry should then be constrained experimentally, by cross-linking mass spectrometry, or by cryo-electron microscopy of the purified proteins, which would test the predicted topology and the arrangement together. Only then is site-directed mutagenesis of specific residues informative: mutating a residue list drawn from a model this uncertain risks a null result that says nothing, because the mutated positions may not be at the real interface.

The residues we report should be treated as a low prior starting list, not as the interface. Second, whether MS4A4A contributes causally to macrophage state should be tested by perturbation: CRISPR knockout and rescue in induced pluripotent stem cell derived or primary human macrophages, and dCas9 based epigenome editing of the promoter region identified here, would separate necessity from correlation, using a targeted activator to test sufficiency [53] and a targeted repressor to test necessity [54]. Beyond these, single-cell multi-omic profiling, with RNA and ATAC from the same cells, would close the gap between our two bulk assays; TREM2-DAP12 signalling assays would test the proposed mechanistic link, using mutants only once the interface has been established independently; and extension to microglia, tumour-associated macrophages, and additional disease contexts would establish how general the association is. Epitope mapping and binding studies for MS4A directed antibodies such as AL044 would connect this work to the therapeutic programme already in the clinic [14].

### 4.6. Therapeutic Implications

MS4A4A’s membrane localization and myeloid-restricted expression make it, in principle, both accessible and selective for antibody-based approaches, and the detection pattern seen at single-cell resolution, a difference in the fraction of detected cells rather than in per-cell level, is the behaviour a marker needs if it is to identify a target population. Context dependency is decisive: M2-like macrophages can be beneficial, in tissue repair and resolution of inflammation, or detrimental, in tumour immunosuppression, so any therapeutic strategy must be framed for a specific disease context rather than as global activation or inhibition of an M2 programme. The substantial donor-to-donor variation observed here in both transcript level and promoter accessibility suggests that patient stratification may be necessary. These data establish associations and generate hypotheses; they do not demonstrate a causal target or a therapeutic window.

## 5. Conclusions

Integrating multi level public data with structure prediction, we find that MS4A4A is elevated in M2 relative to M1 human macrophages at the transcript level, that its promoter shows the highest mean signal in M2 and the lowest between-donor variability among the macrophage states, across three donors, that it is macrophage-enriched in liver and monocyte-enriched in blood with expression concentrated in the M2 leaning cells of each compartment, and that its predicted transmembrane core is a confidently modelled four-helix bundle (mean pLDDT 83.1) attached to long terminal segments the model does not resolve, so that the model as a whole is not confident (62.4). The predicted association with MS4A6A, by contrast, is a hypothesis whose interface the model does not resolve. These results constitute multi level association rather than proof of causation, and a calibrated structural hypothesis rather than structural proof. Stated at that strength, they provide a defensible framework for the experimental work needed to establish whether MS4A4A regulates macrophage state, and whether it is a viable therapeutic target in inflammation and cancer.

## Supplementary Materials

The following supporting information can be downloaded at the journal supplementary material URL, supplied by the editorial office at proof stage. Table S1: Datasets, reference structures, and structure predictions used in this study; Table S2: Complete DESeq2 results for the M2 versus M1 contrast under the donor-blocked model, 12,900 genes (comma separated file); Table S3: MS4A4A and MS4A6A pseudocounts per-sample, by state and donor; Table S4: M2 versus M1 differential expression statistics for MS4A4A and MS4A6A under both models; Table S5: Donor mean normalized ATAC-seq coverage at four named features of the MS4A4A locus; Table S6: MS4A4A promoter accessibility per-donor in four cell states, with coverage completeness; Table S7: MS4A4A detection frequency by group; Table S8: Mean log normalized expression of MS4A4A associated markers in MS4A4A-detected versus negative blood monocytes, with a test per marker; Table S9: Residues at the predicted MS4A4A- MS4A6A interface, with the contact criterion used; Table S10: Complete results for the same contrast without donor blocking, 12,900 genes (comma separated file); Table S11: Normalized

ATAC-seq coverage in bins of 100 bp for 12 samples, 5,040 rows (comma separated file); Table S12: AlphaFold 3 prediction runs and model confidence metrics; Table S13: Software and package versions by figure panel. Supplementary Software S1: a single archive containing all analysis scripts, the input data they read, and the environment records. It comprises the six R scripts and one Python script that generate every computed figure panel and every derived supplementary table; the bulk RNA-seq count matrix and sample table (GSE162698); the liver single-cell count matrix and per cell annotations (GSE115469, 8,444 cells); the blood single-cell count matrix as analysed and its per cell annotations (GSE150728, the 8,006 cell subsample); a data provenance record for each single-cell dataset; the extracted ATAC-seq coverage read by the coverage script; the Figure 1 panel “sessionInfo()” records; and a manifest listing every file with its SHA-256 checksum. The analysis runs end to end from this archive.

## Supporting information

Supplementary tables and codes

## Author Contributions

Shi-Ruei Lin: methodology, software, formal analysis, investigation, data curation, visualization, and writing—original draft preparation. Mandarina Li: methodology, software, formal analysis, investigation, data curation, visualization, and writing—original draft preparation. Sihan Wang: methodology, software, formal analysis, investigation, data curation, visualization, and writing— original draft preparation. Eric Li: methodology, software, formal analysis, investigation, data curation, visualization, and writing—original draft preparation. Hao Sun: conceptualization, validation, resources, supervision, project administration, and writing—review and editing. Liangyu Li: conceptualization, validation, resources, supervision, project administration, and writing—review and editing. All authors have read and agreed to the published version of the manuscript.

## Institutional Review Board Statement

Not applicable. This study analysed only previously published, publicly available, de-identified human data; no new human or animal material was collected.

## Informed Consent Statement

Not applicable. Informed consent was obtained by the investigators of each source study, as reported in the original publications cited in Table S1.

## Data Availability Statement

No new biological samples or sequencing data were generated in this study. This work is a reanalysis of publicly available human datasets. All source data are available in the public domain: Gene Expression Omnibus accessions GSE162698 (bulk RNA-seq; https://www.ncbi.nlm.nih.gov/geo/query/acc.cgi?acc=GSE162698), GSE261697 (bulk ATAC- seq; https://www.ncbi.nlm.nih.gov/geo/query/acc.cgi?acc=GSE261697), GSE115469 (liver single-cell RNA-seq; https://www.ncbi.nlm.nih.gov/geo/query/acc.cgi?acc=GSE115469) and GSE150728 (blood single-cell RNA-seq; https://www.ncbi.nlm.nih.gov/geo/query/acc.cgi?acc=GSE150728); Protein Data Bank entry 6Y90 (https://doi.org/10.2210/pdb6Y90/pdb); and UniProt entries Q96JQ5 and Q9H2W1. All analysis code written for this study and all input data read by that code are submitted with this manuscript as Supplementary Software S1, which also carries a manifest with a SHA-256 checksum for every file; the analysis reported here runs end to end from that archive. The complete derived result tables are provided as Tables S2, S10 and S11, and the remaining derived tables as Tables S3 to S9, S12 and S13. The only input not redistributed is the set of GSE261697 bigWig coverage files, which are available from the accession above and are required only to regenerate Table S11 from raw coverage.

## Acknowledgments

The authors thank the investigators of the four source studies for depositing their data publicly, without which this work would not have been possible. During the preparation of this manuscript, the authors used Claude (Anthropic) for drafting and editing of manuscript text, for generating and debugging analysis and figure code, and for checking reference metadata against Crossref and PubMed. No data were generated, altered, or selected by these tools; every value reported was computed by the analysis scripts supplied in Supplementary Software S1. The authors have reviewed and edited the output and take full responsibility for the content of this publication.

