## Supplementary tables and codes for "Public-Data Reanalysis Links MS4A4A to M2-like Human Myeloid States and Supports a Predicted Four-Pass Transmembrane Fold": R_session_info_by_panel.docx

**R session information, by figure panel**

MS4A4A in human macrophages. Verbatim sessionInfo() output captured after rendering each panel.

**1 Coverage of this record**

Records are held for the four panels of Figure 1, which were produced by scripts 01 and 02. The same R version, 4.6.1, was used throughout, but not the same machine: Figures 1a and 1b were rendered on macOS 27.0 (aarch64-apple-darwin23) and Figures 1c and 1d on Windows 11 (x86_64-w64-mingw32/x64). Each record below states its own platform.

No session record was captured for the Figure 2 panels, which scripts 03 and 04 produced under R 4.6.1 on the macOS machine. The Seurat 5.5.1 and SeuratObject 5.4.0 versions reported in Supplementary Table S13 were read from the loaded namespace list of the Figure 1a record below rather than from a session record of their own, so they are attested indirectly. Figure 3 involves no R: both models were predicted with AlphaFold 3 on the AlphaFold Server and rendered in UCSF ChimeraX 1.12, and the run parameters are in Supplementary Table S12. Figure 4 is drawn, not computed.

**2 Figure 1a**

R version 4.6.1 (2026-06-24)

Platform: aarch64-apple-darwin23

Running under: macOS Golden Gate 27.0

Matrix products: default

BLAS: /System/Library/Frameworks/Accelerate.framework/Versions/A/Frameworks/vecLib.framework/Versions/A/libBLAS.dylib

LAPACK: /Library/Frameworks/R.framework/Versions/4.6/Resources/lib/libRlapack.dylib; LAPACK version 3.12.1

locale:

[1] en_US.UTF-8/en_US.UTF-8/en_US.UTF-8/C/en_US.UTF-8/en_US.UTF-8

time zone: America/Los_Angeles

tzcode source: internal

attached base packages:

[1] stats4	stats	graphics grDevices utils	datasets methods

[8] base

other attached packages:

[1] EnhancedVolcano_1.30.0	ggrepel_0.9.8

[3] ggplot2_4.0.3	DESeq2_1.52.0

[5] SummarizedExperiment_1.42.0 Biobase_2.72.0

[7] MatrixGenerics_1.24.0	matrixStats_1.5.0

[9] GenomicRanges_1.64.0	Seqinfo_1.2.0

[11] IRanges_2.46.0	S4Vectors_0.50.1

[13] BiocGenerics_0.58.1	generics_0.1.4

loaded via a namespace (and not attached):

[1] deldir_2.0-4	pbapply_1.7-4	gridExtra_2.3.1

[4] rlang_1.3.0	magrittr_2.0.5	RcppAnnoy_0.0.23

[7] otel_0.2.0	spatstat.geom_3.8-2	ggridges_0.5.7

[10] compiler_4.6.1	systemfonts_1.3.2	png_0.1-9

[13] vctrs_0.7.3	reshape2_1.4.5	stringr_1.6.0

[16] pkgconfig_2.0.3	fastmap_1.2.0	XVector_0.52.0

[19] labeling_0.4.3	promises_1.5.0	ragg_1.5.2

[22] purrr_1.2.2	jsonlite_2.0.0	goftest_1.2-3

[25] later_1.4.8	DelayedArray_0.38.2	spatstat.utils_3.2-4

[28] BiocParallel_1.46.0	irlba_2.3.7	parallel_4.6.1

[31] cluster_2.1.8.3	R6_2.6.1	ica_1.0-3

[34] spatstat.data_3.1-9	stringi_1.8.7	RColorBrewer_1.1-3

[37] reticulate_1.46.0	spatstat.univar_3.2-0 parallelly_1.48.0

[40] lmtest_0.9-40	scattermore_1.2	Rcpp_1.1.2

[43] tensor_1.5.1	future.apply_1.20.2	zoo_1.9-0

[46] sctransform_0.4.3	httpuv_1.6.17	Matrix_1.7-6

[49] splines_4.6.1	igraph_2.3.3	tidyselect_1.2.1

[52] rstudioapi_0.19.0	abind_1.4-8	spatstat.random_3.5-1

[55] spatstat.explore_3.8-2 codetools_0.2-20	miniUI_0.1.2

[58] listenv_1.0.0	plyr_1.8.9	lattice_0.22-9

[61] tibble_3.3.1	withr_3.0.3	shiny_1.14.0

[64] S7_0.2.2	ROCR_1.0-12	Rtsne_0.17

[67] future_1.75.0	fastDummies_1.7.6	survival_3.8-9

[70] polyclip_1.10-7	fitdistrplus_1.2-6	pillar_1.11.1

[73] Seurat_5.5.1	KernSmooth_2.23-26	plotly_4.12.1

[76] RcppHNSW_0.7.0	sp_2.2-3	scales_1.4.0

[79] globals_0.19.1	xtable_1.8-8	glue_1.8.1

[82] tools_4.6.1	data.table_1.18.4	RSpectra_0.16-2

[85] locfit_1.5-9.12	RANN_2.6.2	dotCall64_1.2

[88] cowplot_1.2.0	grid_4.6.1	tidyr_1.3.2

[91] nlme_3.1-170	patchwork_1.3.2	cli_3.6.6

[94] spatstat.sparse_3.2-0 textshaping_1.0.5	spam_2.11-4

[97] S4Arrays_1.12.0	viridisLite_0.4.3	dplyr_1.2.1

[100] uwot_0.2.4	gtable_0.3.6	digest_0.6.39

[103] progressr_1.0.0	SparseArray_1.12.2	htmlwidgets_1.6.4

[106] SeuratObject_5.4.0	farver_2.1.2	htmltools_0.5.9

[109] lifecycle_1.0.5	httr_1.4.8	mime_0.13

[112] MASS_7.3-66

**3 Figure 1b**

R version 4.6.1 (2026-06-24)

Platform: aarch64-apple-darwin23

Running under: macOS Golden Gate 27.0

Matrix products: default

BLAS: /System/Library/Frameworks/Accelerate.framework/Versions/A/Frameworks/vecLib.framework/Versions/A/libBLAS.dylib

LAPACK: /Library/Frameworks/R.framework/Versions/4.6/Resources/lib/libRlapack.dylib; LAPACK version 3.12.1

locale:

[1] en_US.UTF-8/en_US.UTF-8/en_US.UTF-8/C/en_US.UTF-8/en_US.UTF-8

time zone: America/Los_Angeles

tzcode source: internal

attached base packages:

[1] stats	graphics grDevices utils	datasets methods	base

loaded via a namespace (and not attached):

[1] deldir_2.0-4	pbapply_1.7-4	gridExtra_2.3.1

[4] rlang_1.3.0	magrittr_2.0.5	RcppAnnoy_0.0.23

[7] otel_0.2.0	matrixStats_1.5.0	ggridges_0.5.7

[10] compiler_4.6.1	spatstat.geom_3.8-2	png_0.1-9

[13] vctrs_0.7.3	reshape2_1.4.5	stringr_1.6.0

[16] pkgconfig_2.0.3	fastmap_1.2.0	promises_1.5.0

[19] purrr_1.2.2	jsonlite_2.0.0	goftest_1.2-3

[22] later_1.4.8	spatstat.utils_3.2-4	irlba_2.3.7

[25] parallel_4.6.1	cluster_2.1.8.3	R6_2.6.1

[28] ica_1.0-3	stringi_1.8.7	RColorBrewer_1.1-3

[31] spatstat.data_3.1-9	reticulate_1.46.0	parallelly_1.48.0

[34] spatstat.univar_3.2-0 lmtest_0.9-40	scattermore_1.2

[37] Rcpp_1.1.2	tensor_1.5.1	future.apply_1.20.2

[40] zoo_1.9-0	sctransform_0.4.3	httpuv_1.6.17

[43] Matrix_1.7-6	splines_4.6.1	igraph_2.3.3

[46] tidyselect_1.2.1	rstudioapi_0.19.0	abind_1.4-8

[49] codetools_0.2-20	spatstat.random_3.5-1 miniUI_0.1.2

[52] spatstat.explore_3.8-2 listenv_1.0.0	lattice_0.22-9

[55] tibble_3.3.1	plyr_1.8.9	shiny_1.14.0

[58] S7_0.2.2	ROCR_1.0-12	Rtsne_0.17

[61] future_1.75.0	fastDummies_1.7.6	survival_3.8-9

[64] polyclip_1.10-7	fitdistrplus_1.2-6	pillar_1.11.1

[67] Seurat_5.5.1	KernSmooth_2.23-26	plotly_4.12.1

[70] generics_0.1.4	RcppHNSW_0.7.0	sp_2.2-3

[73] ggplot2_4.0.3	scales_1.4.0	globals_0.19.1

[76] xtable_1.8-8	glue_1.8.1	tools_4.6.1

[79] data.table_1.18.4	RSpectra_0.16-2	RANN_2.6.2

[82] dotCall64_1.2	cowplot_1.2.0	grid_4.6.1

[85] tidyr_1.3.2	nlme_3.1-170	patchwork_1.3.2

[88] cli_3.6.6	spatstat.sparse_3.2-0 spam_2.11-4

[91] viridisLite_0.4.3	dplyr_1.2.1	uwot_0.2.4

[94] gtable_0.3.6	digest_0.6.39	progressr_1.0.0

[97] ggrepel_0.9.8	htmlwidgets_1.6.4	SeuratObject_5.4.0

[100] farver_2.1.2	htmltools_0.5.9	lifecycle_1.0.5

[103] httr_1.4.8	mime_0.13	MASS_7.3-66

**4 Figure 1c and 1d**

R version 4.6.1 (2026-06-24 ucrt)

Platform: x86_64-w64-mingw32/x64

Running under: Windows 11 x64 (build 26200)

Matrix products: default

LAPACK version 3.12.1

locale:

[1] LC_COLLATE=Japanese_Japan.utf8 LC_CTYPE=Japanese_Japan.utf8

[3] LC_MONETARY=Japanese_Japan.utf8 LC_NUMERIC=C

[5] LC_TIME=Japanese_Japan.utf8

time zone: America/Los_Angeles

tzcode source: internal

attached base packages:

[1] stats	graphics grDevices utils	datasets methods	base

other attached packages:

[1] lubridate_1.9.5 forcats_1.0.1	stringr_1.6.0	dplyr_1.2.1

[5] purrr_1.2.2	readr_2.2.0	tidyr_1.3.2	tibble_3.3.1

[9] ggplot2_4.0.3	tidyverse_2.0.0

loaded via a namespace (and not attached):

[1] bit_4.6.0	gtable_0.3.6	crayon_1.5.3

[4] compiler_4.6.1	tidyselect_1.2.1	parallel_4.6.1

[7] scales_1.4.0	R6_2.6.1	labeling_0.4.3

[10] generics_0.1.4	pillar_1.11.1	RColorBrewer_1.1-3

[13] tzdb_0.5.0	rlang_1.3.0	utf8_1.2.6

[16] stringi_1.8.7	S7_0.2.2	bit64_4.8.2

[19] timechange_0.4.0	cli_3.6.6	withr_3.0.3

[22] magrittr_2.0.5	grid_4.6.1	vroom_1.7.1

[25] rstudioapi_0.19.0 hms_1.1.4	lifecycle_1.0.5

[28] vctrs_0.7.3	glue_1.8.1	farver_2.1.2

[31] tools_4.6.1	pkgconfig_2.0.3
