## Supplementary tables and codes for "Public-Data Reanalysis Links MS4A4A to M2-like Human Myeloid States and Supports a Predicted Four-Pass Transmembrane Fold": Supplementary_Tables.docx

Submitted to *Genes*. All tables in this study are supplementary and are numbered

in the order they are first cited in the manuscript. Tables S2, S10, and S11 are

large data files supplied separately as comma separated files; their legends

appear at the end of this document. Each caption is placed above its table, as the

journal requires. The analysis code that produced these tables, and the input

data it reads, are supplied as Supplementary Software S1.

**Table S1. Datasets, reference structures, and structure predictions used in this study.**

| **Assay or resource** | **Accession** | **Source** | **Material** | **Figure** |
| --- | --- | --- | --- | --- |
| Bulk RNA-seq | GSE162698 | Hoppstadter et al., eBioMedicine 2021 | Primary human monocyte derived macrophages; 3 donors; M0/M1/M2(IL-4)/TAM-like; gene level TPM | 1a, 1b |
| Bulk ATAC-seq | GSE261697 | No associated publication; GEO dataset | Primary human monocytes and monocyte derived macrophages; 3 donors (A to C); monocyte/M0/M1/M2(IL-4); bigWig coverage, hg19 | 1c, 1d |
| scRNA-seq (liver) | GSE115469 | MacParland et al., Nat Commun 2018 | Healthy human liver; 5 donors; 8,444 cells; 10x Genomics | 2a to 2c |
| scRNA-seq (blood) | GSE150728 | Wilk et al., Nat Med 2020 | PBMC; 7 severe COVID-19 patients + 6 healthy donors; representative subsample of 8,006 cells from 44,721; Seq-Well | 2d to 2f |
| Protein sequences | Q96JQ5; Q9H2W1 | UniProt Consortium 2025 | Human MS4A4A (239 aa); human MS4A6A (248 aa) | 3a, 3c to 3f |
| Cryo-EM structure | PDB 6Y90 | Kumar et al., Science 2020 | Full length CD20 (MS4A1) homodimer with two rituximab Fab fragments; 3.69 angstroms; chains A/B = CD20 | 3b |
| Structure prediction | AlphaFold 3 (AlphaFold Server) | Generated for this study (see Table S12) | MS4A4A monomer and two chain MS4A4A-MS4A6A complex; run 29 July 2026 and 30 July 2026 | 3a, 3c to 3f |

**Table S3. MS4A4A and MS4A6A pseudocounts per-sample in GSE162698, by state and donor.**

| **Gene** | **State** | **Donor 1** | **Donor 2** | **Donor 3** |
| --- | --- | --- | --- | --- |
| MS4A4A | M0 | 5791 | 4650 | 2682 |
| MS4A4A | M1 | 2380 | 75 | 329 |
| MS4A4A | M2 | 8769 | 2962 | 2061 |
| MS4A4A | TAM | 4444 | 6608 | 4425 |
| MS4A6A | M0 | 16488 | 16926 | 8557 |
| MS4A6A | M1 | 6466 | 118 | 166 |
| MS4A6A | M2 | 32204 | 14948 | 10385 |
| MS4A6A | TAM | 10681 | 25438 | 14027 |

**Table S4. M2 versus M1 differential expression statistics for MS4A4A and MS4A6A under both models.**

| **Gene** | **Model** | **log_2_ FC** | **lfcSE** | **P** | **adjusted P** | **baseMean** | **Fold change** |
| --- | --- | --- | --- | --- | --- | --- | --- |
| MS4A4A | condition + donor (DESeq2 1.52.0) | 2.678 | 0.728 | 0.000237 | 0.00126 | 3712.4 | 6.4 |
| MS4A4A | condition only (pydeseq2 0.5.2) | 2.327 | 0.876 | 0.0079 | 0.0267 | 3712.4 | 5.02 |
| MS4A6A | condition + donor (DESeq2 1.52.0) | 4.034 | 0.934 | 1.56 × 10^-5^ | 0.000121 | 12860.8 | 16.38 |
| MS4A6A | condition only (pydeseq2 0.5.2) | 3.131 | 1.051 | 0.00289 | 0.0118 | 12860.8 | 8.76 |

**Table S5. Donor mean normalized ATAC-seq coverage at four named features of the MS4A4A locus, as fold over genome mean.**

| **Feature** | **Monocyte** | **M0** | **M1** | **M2** |
| --- | --- | --- | --- | --- |
| Promoter apex (TSS, chr11:60,048,014) | 12.05 | 19.11 | 17.89 | 29.74 |
| Intragenic peak 1 (chr11:60,049,900) | 4.37 | 6.94 | 6.62 | 14.49 |
| Intragenic peak 2 (chr11:60,053,000) | 0.71 | 3.09 | 2.34 | 12.18 |
| Downstream peak (chr11:60,078,200) | 5.96 | 21.94 | 16.63 | 29.56 |

**Table S6. MS4A4A promoter accessibility per donor (mean normalized ATAC-seq coverage over hg19 chr11:60,047,000 to 60,049,599), with coverage completeness.**

| **State** | **Donor A** | **Donor B** | **Donor C** | **Mean** | **SD** | **Fold over monocyte mean** | **Promoter bins with no coverage (of 26)** | **Locus bins with no coverage (%)** |
| --- | --- | --- | --- | --- | --- | --- | --- | --- |
| Monocyte | 2.73 | 2.28 | 1.96 | 2.33 | 0.39 | 1.00 | A 12, B 2, C 1 | A 63.1, B 31.2, C 42.6 |
| M0 | 2.67 | 10.59 | 0.73 | 4.66 | 5.23 | 2.00 | A 0, B 0, C 16 | A 18.8, B 37.1, C 54.5 |
| M1 | 7.78 | 2.92 | 2.13 | 4.28 | 3.06 | 1.84 | A 2, B 2, C 2 | A 27.6, B 23.6, C 27.1 |
| M2 | 10.57 | 9.65 | 7.29 | 9.17 | 1.69 | 3.94 | A 0, B 0, C 0 | A 24.5, B 21.9, C 23.6 |

**Table S7. MS4A4A detection frequency by group. Values are descriptive: proportions and tests are computed across cells, not across donors, so they do not constitute donor-level inference (see Section 2.6).**

| **Group** | **Cells (n)** | MS4A4A-detected (at least 1 count) | **Positive (%)** | **Note** |
| --- | --- | --- | --- | --- |
| Liver macrophages, M2-like (Kupffer cell) | 326 | 250 | 76.7 | Wilcoxon rank sum P = 4.53 × 10^-39^ versus M1-like |
| Liver macrophages, M1-like (inflammatory) | 822 | 323 | 39.3 |  |
| Blood monocytes, COVID-19 | 1,560 | 239 | 15.3 | Full atlas 15.4%; descriptive, untested |
| Blood monocytes, Healthy | 531 | 22 | 4.1 | Full atlas 5.0%; descriptive, untested |
| Blood monocytes, all | 2,091 | 261 | 12.5 | Groups compared in Figure 2f |

**Table S8. Mean log-normalized expression of MS4A4A-associated markers in MS4A4A-detected (n = 261) versus MS4A4A-undetected (n = 1,830) blood monocytes. Fold is the ratio of the two means; P is from a two-sided Wilcoxon rank sum test across cells. Values are descriptive: means, fold differences and Wilcoxon P values are computed across cells and treat cells from the same donor as independent, so they are not donor-level inference (see Section 2.6). Of the four markers, only CD163 is displayed as a panel; the four marker comparisons are exploratory and were not adjusted for multiple testing.**

| **Marker** | **Association** | **Mean, positive** | **Mean, negative** | **Fold** | **Above zero, positive (%)** | **Above zero, negative (%)** | **P** | **Panel** |
| --- | --- | --- | --- | --- | --- | --- | --- | --- |
| CD163 | M2, anti inflammatory | 1.170 | 0.828 | 1.41 | 50.6 | 36.9 | 1.1 × 10^-5^ | Figure 2f |
| MERTK | M2, anti inflammatory | 0.167 | 0.066 | 2.53 | 8.8 | 3.3 | 3.1 × 10^-5^ | Value only |
| C1QA | M2, anti inflammatory | 0.124 | 0.032 | 3.92 | 4.6 | 1.4 | 2.0 × 10^-4^ | Value only |
| TNF | M1, pro inflammatory | 0.037 | 0.080 | 0.47 | 1.9 | 4.0 | 9.1 × 10^-2^ | Value only |

**Table S9. Candidate residues at the predicted MS4A4A-MS4A6A interface.**

| **Chain** | **Protein** | **n** | **Candidate interface residues** | **Annotated segments touched** |
| --- | --- | --- | --- | --- |
| A | MS4A4A | 22 | 62, 65, 68, 69, 72, 76, 79, 80, 83, 87, 90, 91, 92, 171, 172, 173, 176, 177, 179, 180, 183, 187 | cytoplasmic, TM1, extracellular, TM4 |
| B | MS4A6A | 24 | 46, 47, 50, 51, 54, 58, 61, 65, 68, 69, 70, 71, 72, 75, 149, 173, 174, 176, 177, 178, 181, 184, 185, 188 | cytoplasmic, TM1, extracellular, TM4 |

**Contact criterion. Residues were selected in UCSF ChimeraX 1.12 as those with any heavy atom within 4 Å of a heavy atom of the opposite chain in the top-ranked predicted model. Because the model has ipTM 0.59, these are candidate contacts from a single low-confidence model and are not validated interaction sites; they should not be used to design mutagenesis without prior experimental evidence of association.**

Chain B residues are given in MS4A6A's own numbering, 1 to 248, not the concatenated numbering 240 to 487 used on the axes of Figure 3f. The contacts lie in the first and fourth transmembrane helices and their flanking extracellular loops in both chains; the second and third helices contribute none.

**Table S12. AlphaFold 3 prediction runs and model confidence metrics.**

| **Attribute** | **MS4A4A monomer** | **MS4A4A-MS4A6A complex** |
| --- | --- | --- |
| Sequences | Q96JQ5 (239 aa) | Q96JQ5 (chain A 1 to 239) + Q9H2W1 (chain B, 240 to 487) |
| Method | AlphaFold 3 / AlphaFold Server | AlphaFold 3 / AlphaFold Server |
| Run date | 2026-07-29 | 2026-07-30 |
| Random seed | 58014515 | 1735480323 |
| Models | 5 | 5 |
| Mean pLDDT, whole model | 62.44 | not recorded per chain |
| Mean pLDDT, TM helices | 83.09 | not recorded per chain |
| pTM | not applicable, single chain | 0.62 |
| ipTM | not applicable, single chain | 0.59 |
| Panels | 3a, 3c | 3d, 3e, 3f |

**Table S13. Software and package versions used, by figure panel.**

| **Software or package** | **Version** | **Used for** |
| --- | --- | --- |
| R | 4.6.1 | all R panels |
| DESeq2 | 1.52.0 | 1a, 1b |
| EnhancedVolcano | 1.30.0 | 1a |
| ggrepel | 0.9.8 | 1a |
| ggplot2 | 4.0.3 | all R panels |
| tidyverse | 2.0.0 | 1c, 1d |
| readr | 2.2.0 | 1c, 1d |
| dplyr | 1.2.1 | 1c, 1d, 2a to 2f |
| tidyr | 1.3.2 | 1c, 1d |
| Seurat | 5.5.1 | 2a to 2f |
| SeuratObject | 5.4.0 | 2a to 2f |
| patchwork | 1.3.2 | figure assembly |
| Matrix | 1.7-6 | 2a to 2f |
| Python | 3 | 1a cross-check, 1c extraction |
| pydeseq2 | 0.5.2 | 1a cross-check |
| AlphaFold | 3 (AlphaFold Server) | 3a, 3c to 3f |
| UCSF ChimeraX | 1.12 | 3a to 3f |

**Legends for the three data files supplied separately**

**Table S2.** Complete DESeq2 results for the M2 versus M1 contrast in GSE162698 under the donor blocked model (`condition + donor`); 12,900 genes. File: `Table_S2_DESeq2_M2_vs_M1_donorblocked.csv`.

**Table S10. Complete results for the same contrast without donor blocking (`condition` only, pydeseq2 0.5.2); 12,900 genes. File: `Table_S10_DESeq2_M2_vs_M1_no_blocking.csv`. Adjusted P values are NA for genes filtered out by pydeseq2's independent-filtering and Cook's-distance outlier steps; this is the expected behaviour of the procedure and those genes were not called in either direction.**

**Table S11.** Normalized ATAC-seq coverage across hg19 chr11:60,040,000 to 60,082,000 in bins of 100 bp for each of 12 samples (GSE261697); 5,040 rows. File: `Table_S11_ATAC_coverage_100bp_bins.csv`.
